# Age-associated changes in knee osteoarthritis, pain-related behaviors, and dorsal root ganglia immunophenotyping of male and female mice

**DOI:** 10.1101/2022.07.07.499172

**Authors:** Terese Geraghty, Alia M. Obeidat, Shingo Ishihara, Matthew J. Wood, Jun Li, Erika Barboza Prado Lopes, Carla R. Scanzello, Timothy M. Griffin, Anne-Marie Malfait, Rachel E. Miller

**Affiliations:** Department of Internal Medicine, Division of Rheumatology, Rush University Medical Center, Chicago, IL, USA; Department of Medicine, Division of Rheumatology, University of Pennsylvania, Philadelphia, PA, USA; Translational Musculoskeletal Research Center, Corp. Michael J. Crescenz VA Medical Center, Philadelphia, PA; Oklahoma Medical Research Foundation, Oklahoma City, OK, USA; OKC Veterans Affairs Medical Center, Oklahoma City, OK, USA

## Abstract

**Objective:** Osteoarthritis (OA) is a leading cause of chronic pain, yet OA pain management remains poor. Age is the strongest predictor of OA development, and mechanisms driving OA pain are unclear. While injury-induced OA models are useful, only a subset of OA is linked to traumatic injury. Here, we aimed to characterize age-associated joint damage, mechanical sensitization, and dorsal root ganglia (DRG) immune phenotypes in mice of both sexes.

**Methods:** Male or female mice aged 6- or 20-months old were evaluated for histopathologic knee OA, pain-related behaviors, and L3-L5 dorsal root ganglia (DRG) immune characterization via flow cytometry. DRG gene expression in aged mice and humans was also examined.

**Results:** Twenty-month old male mice had worse cartilage degeneration than 6-month old mice. Older female knees showed increased cartilage degeneration, but to a lesser degree than males. Older mice of both sexes had worse mechanical allodynia, knee hyperalgesia, and grip strength compared to younger mice. For both sexes, DRGs from older mice showed decreased CD45+ cells, and a significant increase in F4/80+ macrophages and CD11c+ dendritic cells. Older male DRGs showed increased expression of *Ccl2* and *Ccl5* and older female DRGs showed increased *Cxcr4* and *Ccl*3 compared to 6-month DRGs, among other differentially expresssed genes. Human DRG analysis from six individuals >80 years old revealed elevated *CCL2* in male DRGs compared to females, whereas *CCL3* was higher in female DRGs.

**Conclusions:** Here we show that aging in male and female mice is accompanied by mild knee OA, mechanical sensitization, and changes to immune cell populations in the DRG, suggesting novel avenues for development of analgesic therapies.

## Introduction

Risk factors for osteoarthritis (OA), the most prevalent joint disease, include age, female sex, obesity, and joint trauma (1). Age is one of the strongest predictors of OA development (1, 2), and after the age of 50, radiographic joint changes and joint pain become more common (3). In fact, it is estimated only 1 in 8 persons experiencing symptomatic knee OA are under the age of 45 (4). It has been well studied that aging is associated with cartilage changes that contribute to the development of OA, including, but not limited to reduced cartilage thickness (5), increased proteolytic activity (6), and cellular senescence with abnormal secretory profiles (7–9). However, less is known about the driving mechanisms that promote chronic OA pain and how the aging process is involved. Chronic pain is the main reason OA patients seek medical care, yet therapies to address OA pain are inadequate. Existing drugs, such as non-steroidal anti-inflammatory drugs (NSAIDs) often come with serious side effects when used chronically, especially in an older population (10). In addition, studies have found weak correlations between radiographic knee OA and pain severity (11–13), but imaging of soft tissues showed that synovitis and bone marrow lesions correlated with pain (14, 15). Therefore, there is a need for a deeper understanding of mechanisms that drive joint pain associated with aging in order to identify new analgesic targets.

Both peripheral (16, 17) and central (18) sensitization contribute to chronic pain development (19, 20). The cell bodies of specialized pain-sensing neurons called nociceptors reside in the dorsal root ganglia (DRG). Nociceptive neurons extend an axon to the periphery, *e.g.*, the joint, and another to the dorsal horn of the spinal cord. From the DRG, the pain signal is transmitted to the dorsal horn, where the first synapse occurs, then second-order neurons project to supraspinal regions, and the signal is relayed to higher regions of the brain, where pain perception occurs (reviewed in (21, 22)). Peripheral sensitization is influenced by neuroinflammation in the DRG (23–25), and while an important role for immune cells in modifying pain sensitization has been suggested (26–28), very little is known in the context of age-associated OA. Behavior changes in older mice have been previously demonstrated (29), where the authors found trends in decreased locomotor activity, motor function, acoustic startle response, social behavior, and depression-related behavior in mice by the age of 6-7 months that became more pronounced by 8-12 months of age (29). In the monoiodoacetate (MIA) model of OA, differences in the development of referred mechanical allodynia and microglial responses in the spinal cord have been reported in young *vs.* old mice, suggesting potentially different sensitization mechanisms with age (30).

Most experimental animal studies of OA use surgical or chemical methods to cause OA in young animals (reviewed in (31)). These models offer many benefits, including controlling variability and low cost, but they may not precisely represent the pathological mechanisms involved in OA associated with aging. Therefore, studying naturally occurring OA in aged animals may uncover different mechanisms contributing to progression of OA joint damage and associated pain. Here, we aimed to analyze aging-associated OA characteristics of 6-month and 20-month old male and female C57BL/6 mice by evaluating the knee joint, pain-related behaviors, and phenotypes of immune cells in DRGs. We chose 6-month and 20-month old mice because a 6-month old mouse roughly translates to a mature adult human (approximately 30-40 years old), while a 20-month old mouse correlates with an elderly human (approximately 60-70 years old) (32, 33).

## Methods

### Animals

Wildtype (WT) C57BL/6 male mice were bred in house and aged to either 6 months (n=10) or 20 months old (n=12). WT C57BL/6 male mice aged 10 weeks (n=10) were purchased from Jackson Laboratory. WT C57BL/6 female mice aged 10 weeks (n=10), 6 months (n=6) or 20 months (n=6) were purchased from Jackson Laboratory. In each age group, each mouse was subjected to pain behavior testing, joint histology, and DRG and peripheral blood flow cytometry with the exception of 10-week old animals that only were evaluated for pain behavior testing as a young animal baseline. Animals housed at Rush had unrestricted access to food and water and were kept on a 12-hour light cycle.

For DRG gene expression analyses, 6-, 12-, 18- and 24-month-old male and female C57BL/6 mice were obtained from the National Institute on Aging (NIA) Aged Rodent Colony and housed at Oklahoma Medical Research Foundation (OMRF) for a minimum of one week with unrestricted access to food and water prior to euthanasia and tissue collection. All animal experiments were approved by the respective Institutional Animal Care and Use Committees (IACUC).

### Pain-dependent behaviors

Independent sets of mice were evaluated at 10 weeks (n=10 for both males and females), 6 months (n=10 males, n=6 females), or 20 months (n=12 males, n=6 females) for the following pain behaviors: mechanical allodynia, knee hyperalgesia, and grip strength. Mechanical allodynia in the hind paw was measured by von Frey testing as previously described (34, 35). Knee hyperalgesia was measured by pressure application measurement (PAM) testing as previously described (36), and grip strength testing was done on a square grid device using a grip force meter and normalized to mouse body weight, as previously described (37). Only one behavioral test was performed per day, and animals were allowed to rest at least one day between different tests. Blinding was not feasible as the mice were tested as they came of age.

### Histology

Following pain behavior testing, 6-month (n=10) and 20-month old (n=12) male and 6-month (n=6) and 20-month (n=6) female mice were sacrificed to collect right-side knees for histologic analysis, and peripheral blood and DRGs were collected for flow cytometry as described below. The knees were fixed in 10% natural buffered formalin, then decalcified in EDTA for 3 weeks, processed routinely and embedded in paraffin. The knees were serially sectioned at six-micron thickness in the coronal plane. Sections from the center of the joint were stained with Safranin O/Fast Green for the evaluation of cartilage damage. Briefly, 0.1% solution Safranin O and Fast Green solutions were applied to sections for three minutes each at room temperature. Sections were imaged on a light microscope and examined for cartilage degeneration based on Osteoarthritis Research Society International recommendations, as previously described (38, 39). We also measured osteophyte width via an ocular micrometer if osteophytes were present. The synovial pathology was evaluated in the mouse knees as follows: Changes in synovial hyperplasia, cellularity and fibrosis were evaluated at the synovial insertion of lateral femur, medial femur, lateral tibia, and medial tibia separately as described (40, 41). Synovitis scoring was performed by 2 independent observers blinded to the groups. Total limb score of the 3 parameters (synovial hyperplasia defined as thickness of the lining layer with a score range of 0-3, cellularity defined as the cell density of the synovial sublining with a score range of 0-3, and fibrosis defined as the extracellular matrix density in the synovium with a score range of 0-1) at all 4 quadrants was averaged for the two readers and reported per knee.

### Flow cytometry

To yield at least 1 million cells for flow cytometry, 6 DRGs needed to be collected. For males, DRGs from 2 mice were pooled such that lumbar level L3-L5 DRGs of either the right or left side from two mice were collected (3 DRGs per mouse = total 6 DRGs), *i.e.*, when the mouse number n=10, flow cytometry sample number n=5. Trends were similar on right and left sides, thus, flow cytometry plots show right side only data for males. Since we did not see differences between right and left side DRGs in males, we aimed to reduce the number of mice needed per sample for females (*i.e.*, one mouse instead of two mice per sample). Thus, for females, we pooled right and left side L3-L5 DRGs together for one mouse per sample (n=6). After dissection of the L3-L5 knee-innervating DRGs, tissue was digested with collagenase type IV (final concentration = 1.6mg/mL) and DNase I (final concentration = 200ug/mL) for 1 hour shaking at 37°C. Following digestion, cells were counted, and 1 million cells were stained with an immune cell panel of anti-mouse antibodies: PE-CD45, AF700-CD3, BV711-CD11b, PE/Cy7-MHCII, PerCP/Cy5.5-Ly6G, BV605-Ly6C, APC-F4/80, PE/Dazzle-CD206, BV650-CD11c, BV785-CX3CR1, BV421-CD163 (BioLegend), and Aqua-Live/Dead stain (ThermoFisher). After staining, sample data were acquired through an LSR Fortessa flow cytometer. Flow cytometry analysis was completed with FlowJo software (version 10). A representative gating strategy is shown in Suppl. Figs. 1,2 (DRG) and Suppl. Fig. 3 (Peripheral blood).

### Mouse DRG qPCR analysis

Immediately following euthanasia, L3-L5 DRGs were harvested from the left and right side, immediately placed in TRIzol® Reagent (Ambion), and stored at −80°C until homogenization. Samples were mechanically homogenized in TRIzol, and RNA was isolated using RNeasy mini kit (Qiagen) following manufacturer instructions. 224 ng RNA was reverse transcribed to cDNA, pre-amplified, and loaded at 1:20 dilution into a custom Fluidigm DELTAgene Assay 96.96 Dynamic Array IFC for automated PCR reactions run on BioMark HD (Fluidigm), with 88 genes assessed per sample. Standard QC steps were applied using melting curve analysis, reference gene normalization, and sample amplification. For male mice, 31 samples passed QC evaluation with n=8 for 6-month, n=8 for 12-month, n=7 for 18-month, and n=8 for 24-month old mice. For female mice, 30 samples passed QC evaluation with n=6 for 6-month and n=8 for 12-, 18-, and 24-month old mice.

### Human DRG in situ hybridization

Human DRGs came from participants in the Religious Orders Study (ROS) or Rush Memory and Aging Project (MAP) (42, 43). At enrollment, participants agreed to annual clinical evaluation and organ donation at death, including brain, spinal cord, nerve, and muscle. Both studies were approved by an Institutional Review Board of Rush University Medical Center. All participants signed an informed consent, Anatomic Gift Act, and a repository consent to allow their resources to be shared. The DRGs were removed postmortem and flash frozen as part of the spinal cord removal. Clinical characteristics of the human DRGs can be found in **Table 1**.

**Table 1.** Human DRG clinical characteristics from the ROSMAP study used for RNA analysis.

| Sex | Age at death (Years) | BMI |
| --- | --- | --- |
| Female | 86.5 | 27.6 |
| **Female** | **96.2** | **23.0** |
| Female | 89.2 | 27.8 |
| **Male** | **82.0** | **21.3** |
| Male | 93.5 | 24.4 |
| **Male** | **97.8** | **25.8** |
| **Average** | **90.9** | **25.0** |

The human DRGs were embedded in optimal cutting temperature (OCT) compound. Tissues were cryo-sectioned onto slides (20 μm) and frozen until use. RNAscope Multiplex Fluorescent V2 Assay was performed according to Advanced Cell Diagnostics (ACD) Bio instructions with minor modifications, detailed below. Targets included human *CCL2* (423811) and *CCL3* (455331). Slides were fixed in 4% paraformaldehyde (PFA) for 30 min, followed by dehydration through 5 min of 50%, 75% and two 100% ethanol washes. Hydrogen peroxide (3%) was applied for 10 min, and target retrieval performed with a reduced time of 3 min. Protease III was applied for 30 min. After washing, sections were incubated with probes for *CCL2* and *CCL3* at 1:50 dilution at 40°C for 2 hours to hybridize. The signal was amplified, and opal dyes (Akoya Biosciences) were added to detect respective probes at 1:100 dilution. Slides were counterstained with Vectashield containing DAPI. Tissue slides were imaged on a Fluoview FV10i confocal microscope. Laser intensity for all samples was ≤9.9%. Single planes of focus were selected and processed using Fiji software, with only brightness and contrast tools used to adjust images. Quantification was performed manually in a blinded manner by counting the positive signal (number of dots per area) in five different areas per section at 60x magnification.

### Statistical analysis

Sample sizes were chosen based on published data (44) – we calculated that to achieve 80% power with α = 0.05, n > 5 mice would be needed to detect changes in cartilage pathology in male C57BL/6 mice from age 6 months to age 20 months. Statistical analysis by unpaired two-tailed Student’s t test was used for pairwise comparisons, except for synovitis scoring where Mann-Whitney test was used given non-normal distribution of data. Ordinary one-way ANOVA was used for comparison of multiple groups with Tukey post test. For mechanical allodynia, since the von Frey fibers are on a log scale, the data was first log-transformed and then ordinary one-way ANOVA test was done with Tukey post test. Mean +/- standard error of the mean (SEM) is shown in all graphs and p values are stated on each graph. P values were considered significant if less than 0.05. Statistical calculations were performed using GraphPad Prism 9.

## Results

### Knee osteoarthritis in aged mice

We first aimed to assess whether natural aging was associated with histopathologic development of OA. Knees from male and female mice showed no cartilage degeneration in either compartment at the age of 6 months. By 20 months, male mice had significant cartilage damage, both in the medial and lateral compartments (Fig.1A-C), consistent with previous literature (44). Female mice showed significant cartilage degeneration on both the medial and lateral sides from 6 to 20 months, but to a much lesser degree than male mice (Fig. 1F-H). In addition, the knees of 20-month old male mice showed small osteophytes (less than 100 µm in width), while no osteophytes were present in 6-month old male mice. Knees from female mice showed small chondrophytes/osteophytes at both 6 months and 20 months (less than 50 µm in width). Representative images per group (*i.e.*, 6-month, 20-month, male or female knees) are shown in Suppl. Fig. 4. To show the wide range of cartilage degeneration seen in male mice, we show images from 20-month old male mice that have low-to-high degeneration scores on either the medial or lateral sides in Suppl. Fig. 5.

**Fig. 1.**
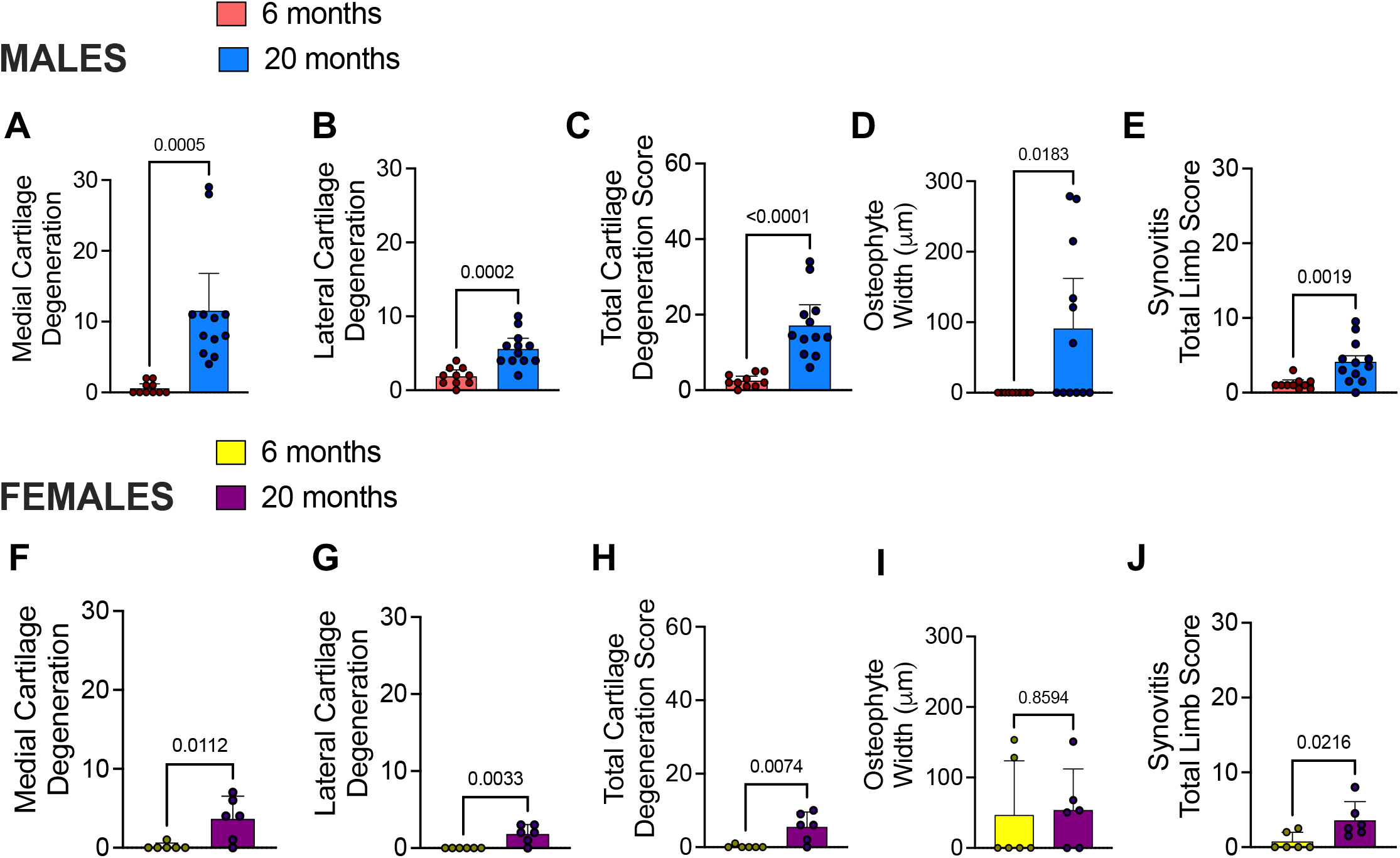
Aging is associated with mild knee osteoarthritis in mice. ***(A)*** Medial cartilage degeneration scores for male mice right knees aged 6 months (n=10) or 20 months (n=12) old as determined by OARSI scoring methods; ***(B)*** lateral cartilage degeneration; ***(C)*** Total cartilage degeneration score for males plotted as sum of medial and lateral compartments; ***(D)*** Osteophyte width for males plotted in micron size found in either medial or lateral compartments of knee ***(E)*** Synovitis total limb score for males mice showing an average of two blinded scores (total limb score sum accounts for synovium hyperplasia, cellularity, and fibrosis – maximum score is 28); ***(F – J)*** Same as in *(A – E)* but shown for female mice aged 6 or 20 months old (n=6). Statistical analysis by unpaired two-tailed t-test, except for Synovitis score where Mann-Whitney test was used. Significant if p < 0.05. Error bars show Mean +/- SEM.

No synovial pathology was detected in either male or female mice at the age of 6 months. However, older mice of both sexes showed mild synovial pathology at 20 months (Fig. 1E+J). Both male and female mice showed hyperplasia in the lining layer of synovium compared to younger mice (Suppl Fig. 6 A+D). No significant changes were detected in cell density of the subintimal layer of male and female mice, yet a trend of increased cellularity was detected in old female mice (Suppl Fig. 6 B+E). In addition, older female mice showed fibrotic subintimal synovium compared to younger counterparts (Suppl Fig. 6 C+F). Overall, these results suggest mild joint degeneration occurs naturally with aging.

### Pain-related behaviors in mice with aging-associated osteoarthritis

We evaluated behaviors indicative of mechanical sensitization in 10-week, 6-month, or 20-month old mice. Mechanical allodynia of the hind paw progressively developed with age in both sexes (shown in Fig. 2A+D). Assessment of knee hyperalgesia revealed no difference in knee withdrawal threshold between 10-week old and 6-month old mice, but knee hyperalgesia developed by 20 months of age compared to 6-month old mice, again in both sexes (shown in Fig. 2B+E). These results suggest aging and naturally occurring knee joint damage is accompanied by increased mechanical sensitization in the hind limb. Finally, grip strength testing revealed a progressive decline in grip strength with age in both sexes, similar to the trend seen in the hind paw mechanical allodynia assay (shown in Fig. 2C+F). Grip strength data was normalized to mouse bodyweights, which are shown in Suppl. Fig. 7.

**Fig. 2.**
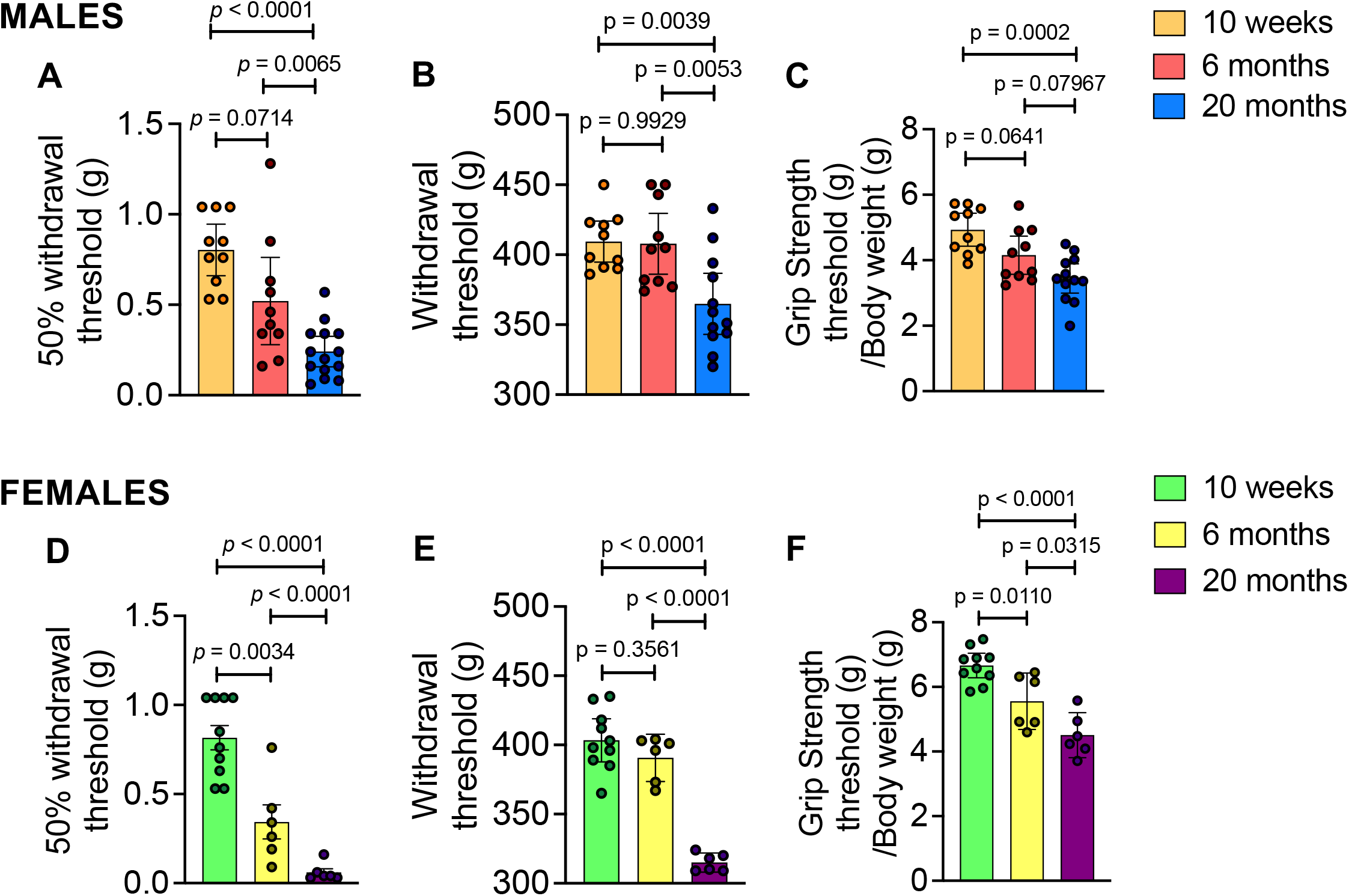
Aging is associated with development of pain-related behaviors. ***(A)*** Mechanical allodynia by von Frey fiber testing ***(B)*** Knee hyperalgesia by pressure application measurement testing and ***(C)*** Grip strength by square grid grip testing of all four paws observed in 10-week (n=10), 6-month (n=10), or 20-month (n=12) old male mice. ***(D - F)*** Same as in *(A - C)* but for 10-week (n=10), 6-month (n=6), or 20-month (n=6) old females. Statistical analysis by Ordinary One-Way ANOVA test. Mechanical allodynia values were first log transformed before running statistical analysis as the von Frey fibers are on a log scale. Significant if p < 0.05. Error bars show Mean +/- SEM.

### DRG immune cell characterization of aged mice

Flow cytometry of DRGs from 20-month old mice showed changes in a number of cell types compared to younger (6 month) mice in both males and females. In particular, 20-month old mice had a decreased total relative frequency of CD45+ cells (significant in males shown in Fig. 3A and trending in females shown in Fig. 4A). In contrast, an increase in F4/80+ macrophages (significant in males shown in Fig. 3B and trending in females shown in Fig. 4B) and a significant increase in CD11c+ dendritic cells in both males and females (Figs. 3C and 4C) was observed in 20-month old mice compared to younger (6 month) mice. For both sexes, there were no significant differences in relative frequencies of other cell types, including CD3+ T cells (Figs. 3D and 4D), CD11b+ myeloid cells (Figs. 3E and 4E), Ly6Chi monocytes (Figs. 3F and 4F), and Ly6G+ granulocytes (Figs. 3G and 4G). In females, there was no difference in proportion of Ly6C-antigen presenting precursor cells between DRGs from 20-month old mice compared to 6-month old mice (Fig. 4H), but this population significantly increased in DRGs from older males (Fig. 3H). Additionally, there was no significant difference in the frequency of CCR2+ cells between younger and older male DRGs (Fig. 3I); however, there was a significant decrease in the relative frequency of CCR2+ cells in DRGs from 20-month old female mice compared to 6-month old female mice (Fig. 4I).

**Fig. 3.**
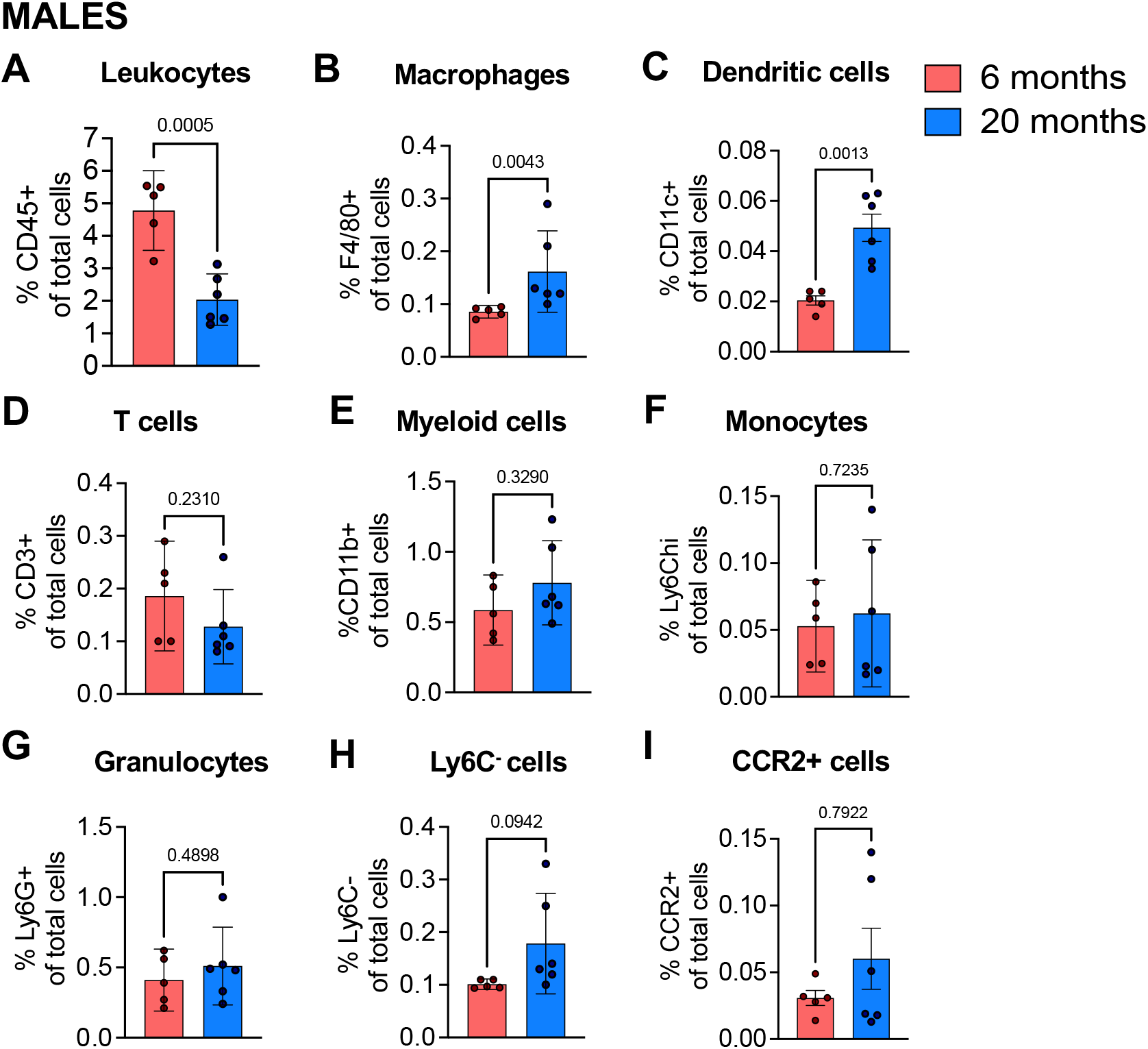
Immune cell populations in dorsal root ganglia of aged male mice. ***(A)*** Frequency of CD45+ leukocytes ***(B)*** CD11b+ myeloid cells ***(C)*** CD3+ T lymphocytes ***(D)*** Ly6Chi monocytes ***(E)*** Ly6G+ granulocytes ***(F)*** CD11c+ dendritic cells ***(G)*** Ly6C-antigen presenting precursor cells ***(H)*** CCR2+ cells ***(I)*** F4/80+ macrophages in 6- month (n=5), or 20-month (n=6) old male mice. Flow cytometry gating strategy in Suppl. Fig. 2. Statistical analysis by two-tailed student’s t-test. Significant if p < 0.05. Error bars show Mean +/- SEM.

**Fig. 4.**
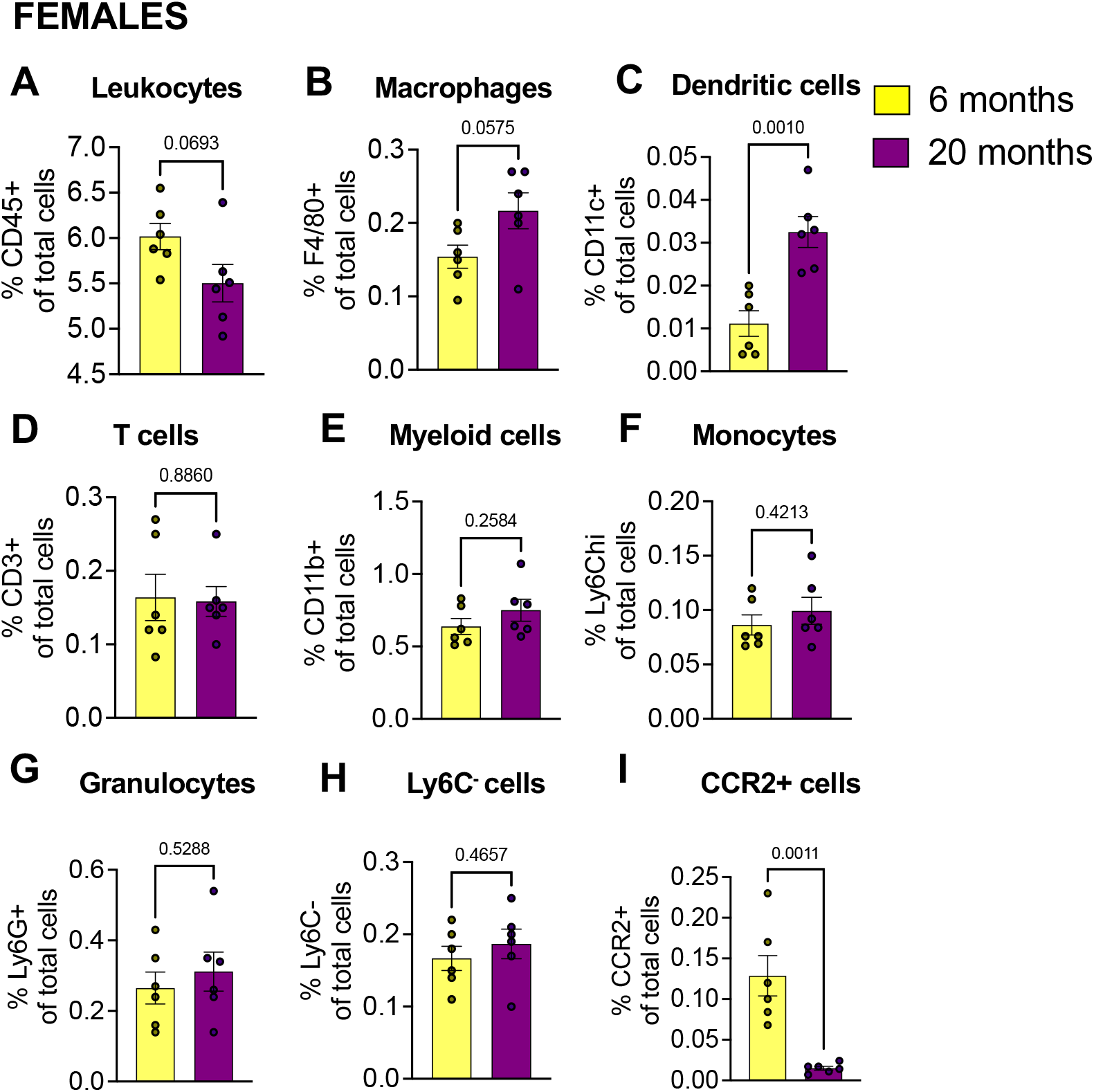
Immune cell populations in dorsal root ganglia of aged female mice. ***(A)*** Frequency of CD45+ leukocytes ***(B)*** CD11b+ myeloid cells ***(C)*** CD3+ T lymphocytes ***(D)*** Ly6Chi monocytes ***(E)*** Ly6G+ granulocytes ***(F)*** CD11c+ dendritic cells ***(G)*** Ly6C-antigen presenting precursor cells ***(H)*** CCR2+ cells ***(I)*** F4/80+ macrophages in 6-month (n=6), or 20-month (n=6) old female mice. Flow cytometry gating strategy in Suppl. Fig. 2. Statistical analysis by two-tailed student’s t-test. Significant if p < 0.05. Error bars show Mean +/- SEM.

We also evaluated macrophage subtypes within the DRG to assess macrophage activation and characterize if they are more pro- or anti-inflammatory. To discriminate between these functional phenotypes, we stained for CD163 and CD206 (also known as mannose-C-type lectin receptor-1 (MRC1)), scavenger receptors associated with an anti-inflammatory phenotype, and CX3CR1 and MHCII, receptors associated with an activated, pro-inflammatory phenotype (45, 46). We found a significant increase in CX3CR1+, MHCII+, and CD206+ macrophages, but no change in CD163+ macrophages in male mice at 20 months compared to 6 months (Fig. 5A-D). There were no significant changes in the number of these macrophage subtypes in DRGs from female aged mice (Fig. 5E-H). Representative gating strategies depicting how this data was acquired and analyzed are shown in Suppl. Figs. 1 **and** 2.

**Fig. 5.**
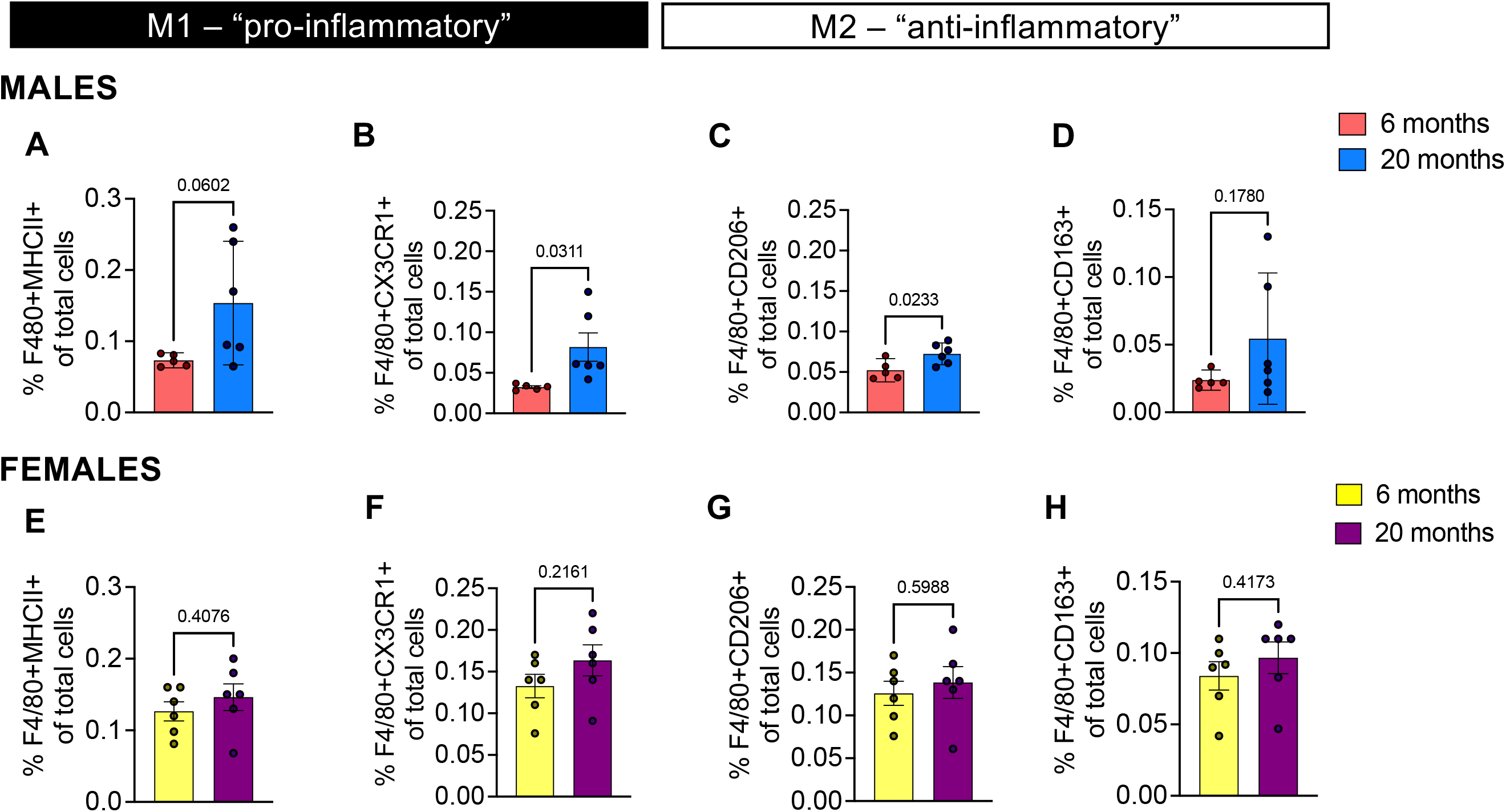
Macrophage phenotypes in male and female aged DRGs. ***(A)*** Frequency of F4/80+MHCII+ pro-inflammatory macrophages ***(B)*** F4/80+CX3CR1+ pro-inflammatory macrophages ***(C)*** F4/80+CD206+ anti-inflammatory macrophages ***(D)*** F4/80+CD163+ anti-inflammatory macrophages from 6-month (n=5) or 20-month old (n=6) male mice DRGs ***(E - H)*** Same as in *(A - D)* but for 6-month or 20-month old female mice (n=6) DRGs. Flow cytometry gating strategy in Suppl. Fig. 3. Statistical analysis by two-tailed student’s t-test. Significant if p < 0.05. Error bars show Mean +/- SEM.

Additionally, to assess any systemic immune changes in naturally aged mice, we examined the circulating immune cells in peripheral blood of aged mice. In both males and females there was a significant decrease in total CD45+ leukocytes in 20-month-old blood compared to 6-month-old blood. There were no significant changes in myeloid cells, granulocytes, monocytes, Ly6Clo or Ly6C-cells in circulation, for both sexes (Suppl. Figs. 8A-F **and** 8J-N), except a slight increase in Ly6C-cells circulating in male mice (Suppl. Fig. 8O). There were no changes in T cell populations in females (Suppl. Figs. 8G-I), but significantly decreased frequencies of CD3+, CD4+, and CD8+ T cells in peripheral blood of male mice (Suppl. Fig. 8P-R). This is consistent with reported literature where it has been reported that peripheral blood CD4+ T cells and CD8+ T cells declined in male C57BL/6 mice by the age of 20-26 months (47). A representative gating strategy for peripheral blood flow cytometry depicting how this data was acquired and analyzed is shown in Suppl. Figs. 3.

### Gene expression analysis of aged mice L3-L5 DRGs

To assess DRG immune changes in aged mice on a molecular level, RNA was extracted from DRGs collected from mice of different ages (6, 12, 18, and 24 months). There were 88 genes assessed and results are shown in Table 2 (males) and Table 3 (females). In male mice, there was significantly increased mRNA expression of *Ccl5, Tacr1, Ccl2, Spp1, Tnf, Tgfb1, and Ccl3* and significantly decreased *Gss* gene expression at 24 months compared to 6 months (**Table 2**). In females, *Cxcr4*, *Hgf, Tacr1, Il1rn, Runx3, Scn9a, Ccl3*, *Kit, Mrgprd, Cx3cr1, Ptk2b, Trpv1, Ntrk1, Trpa1, Tnf, Gclc, Gsr, Npy, Scn10a, Calca, Etv1*, and *P2rx3* were significantly increased at 24 months compared to 6 months, while *Gstsa4, Prkcq*, and *Gstm1* gene expression were decreased (**Table 3**). These data suggest a role for chemokine signaling and immune cell communication within the DRG in aging-associated spontaneous OA.

**Table 2.**
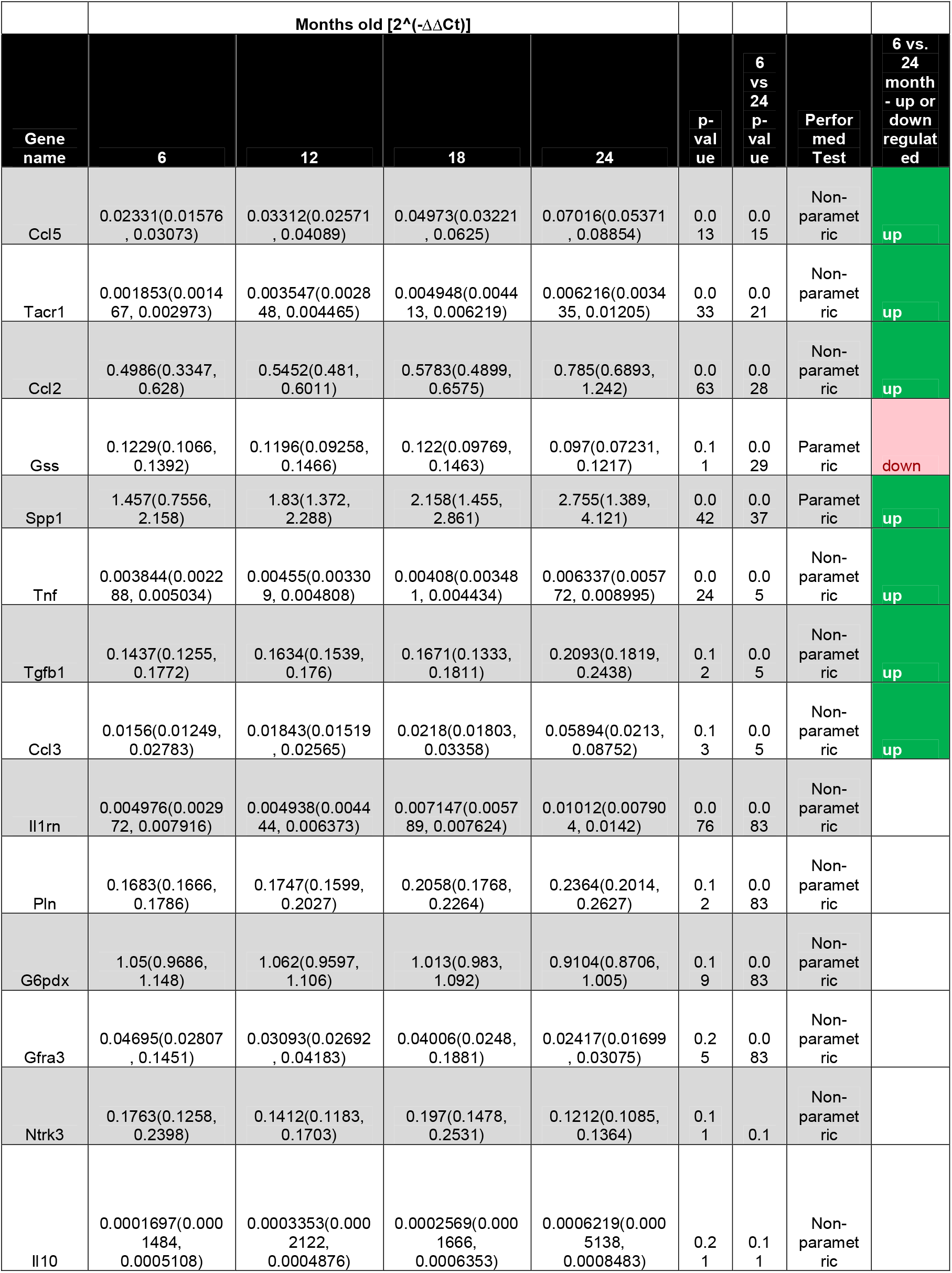
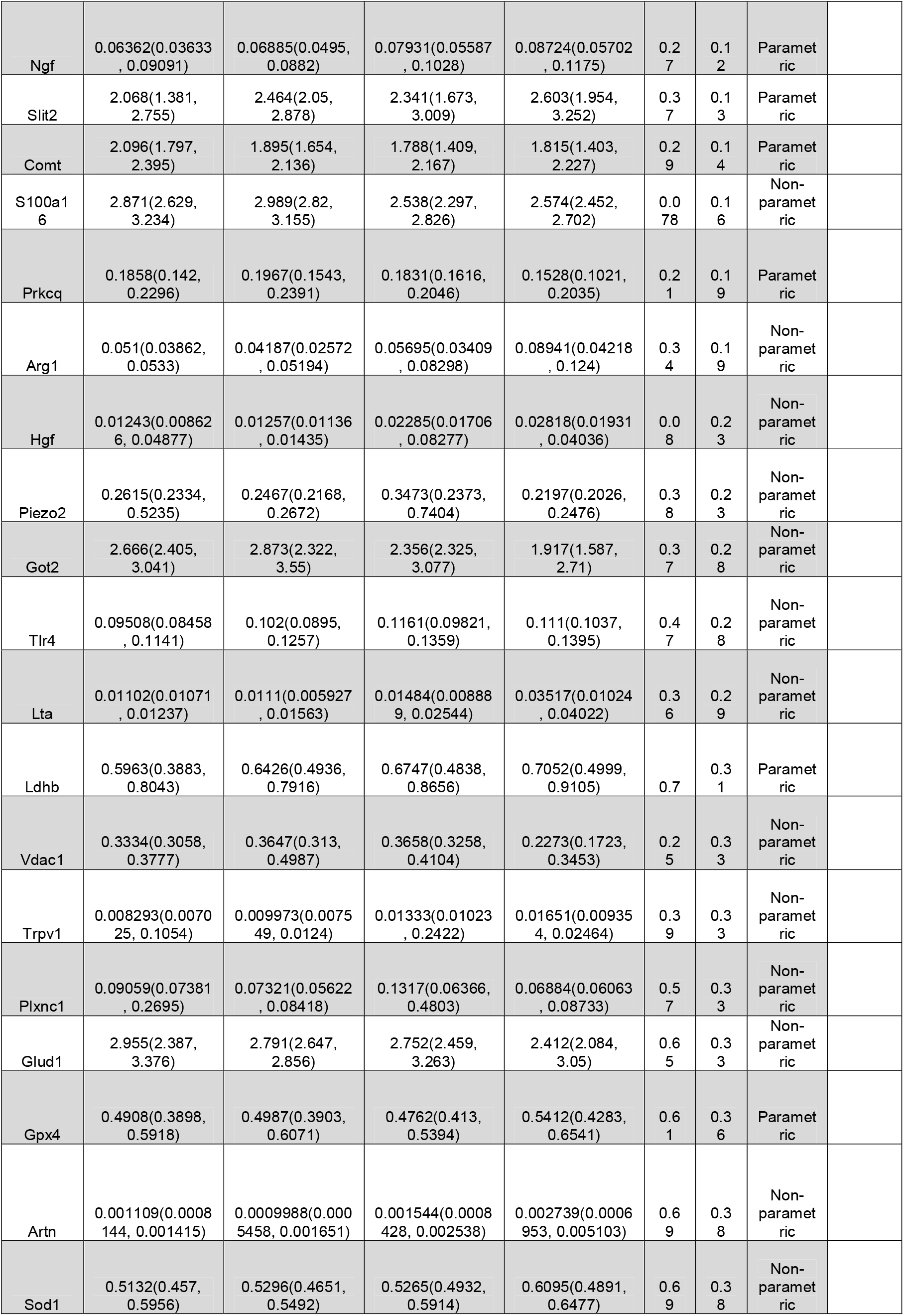
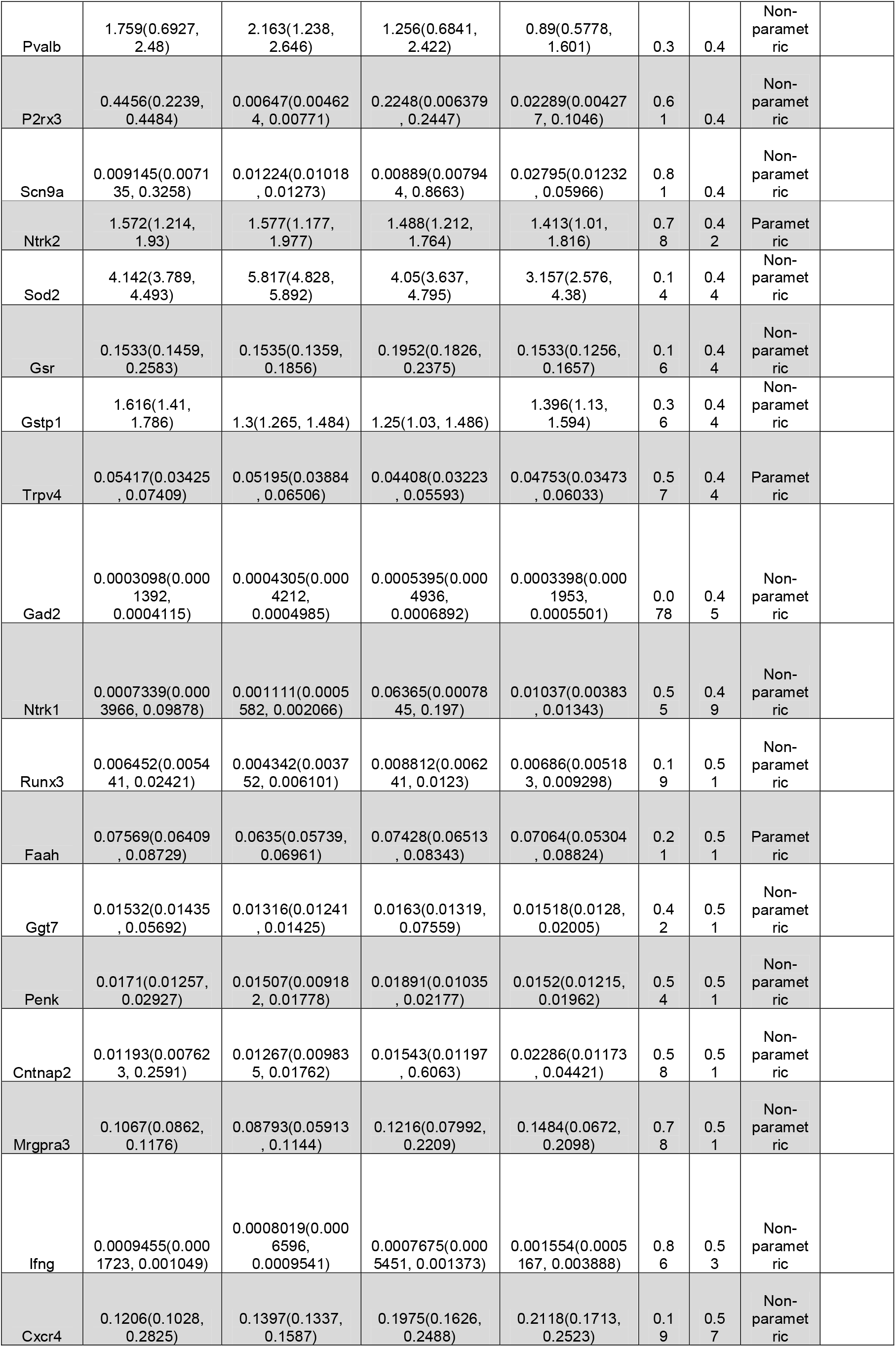
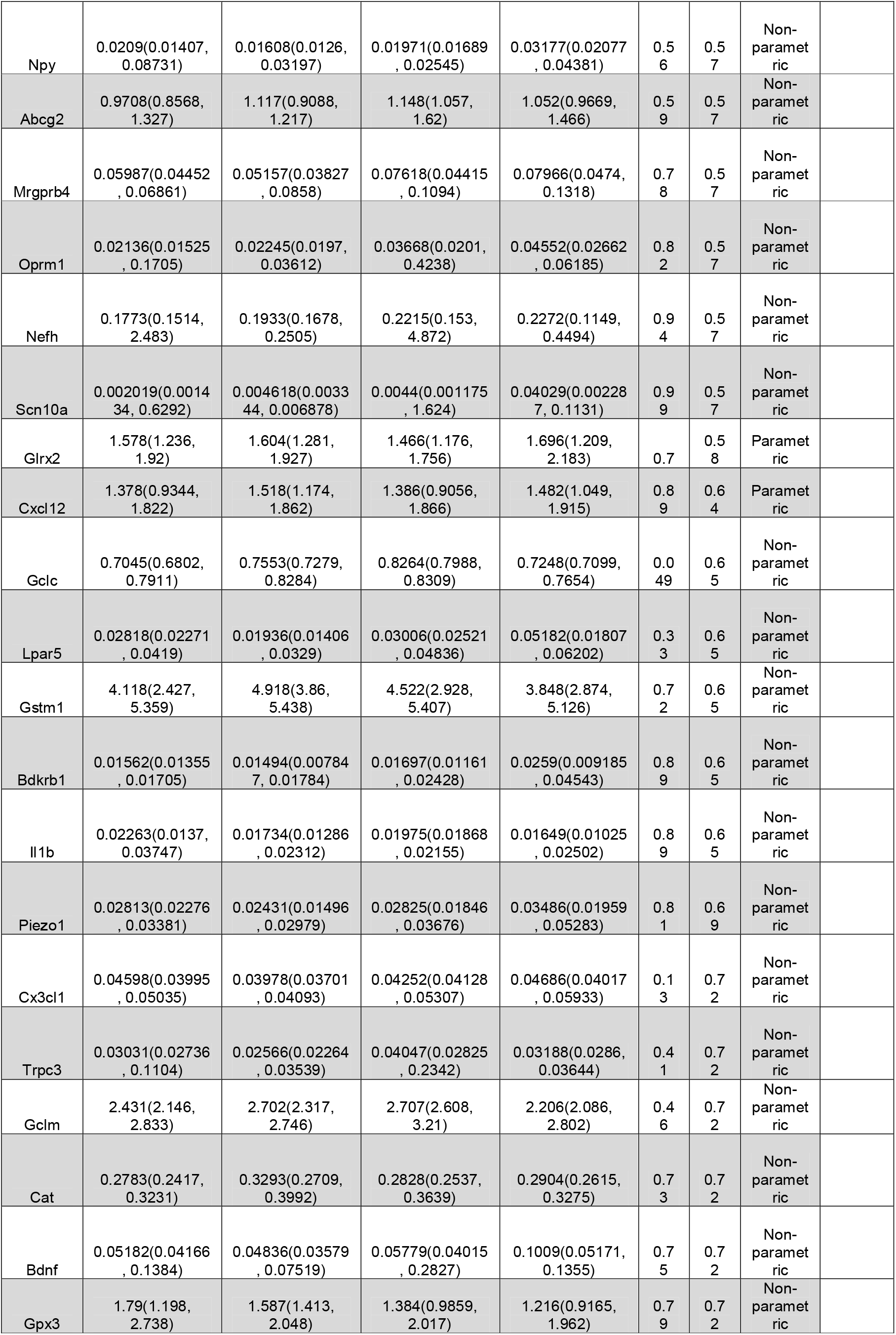
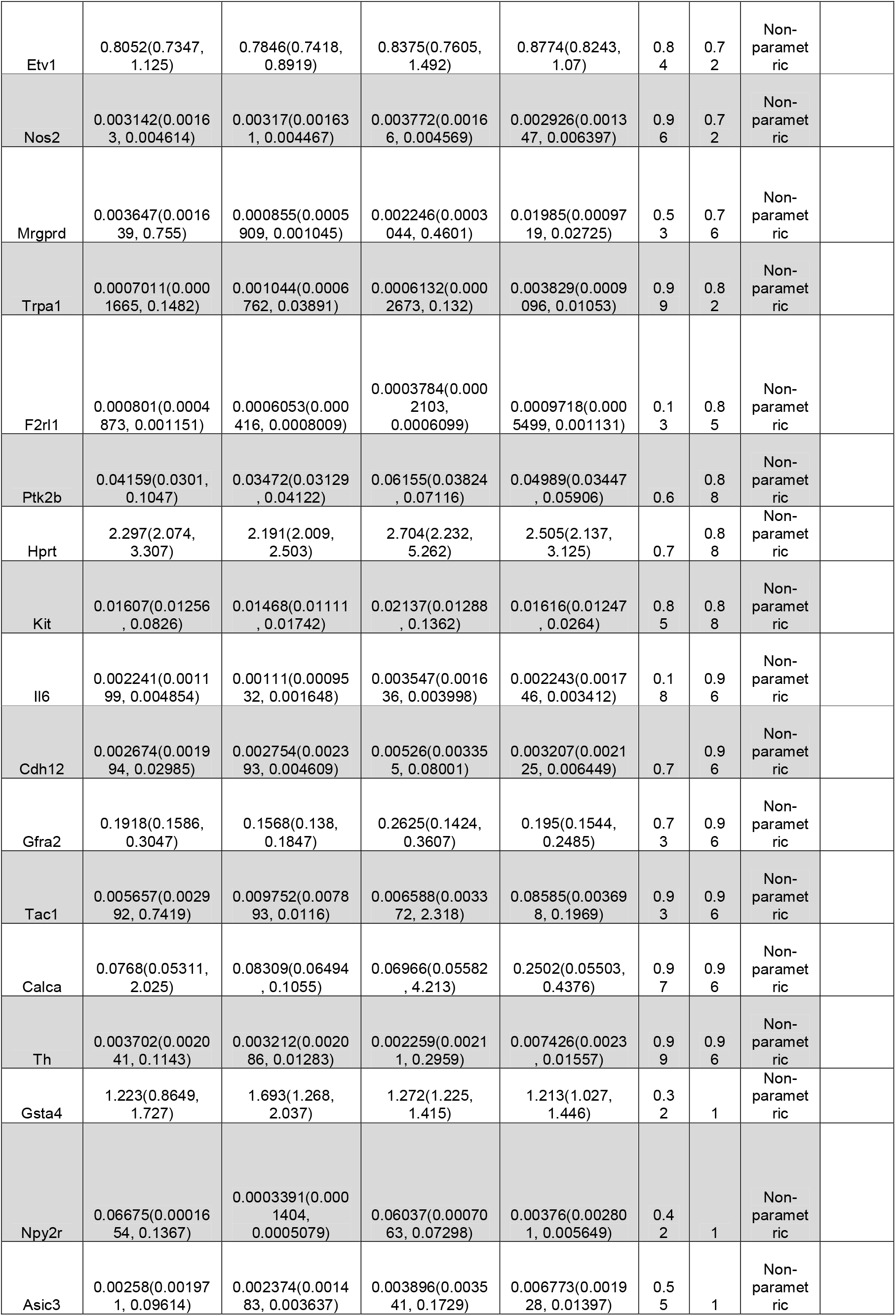
Gene expression in DRGs of male C57BL/6 mice aged 6-months (n=8), 12-months (n=8), 18-months (n=7), and 24-months (n=8) old.

**Table 3.**
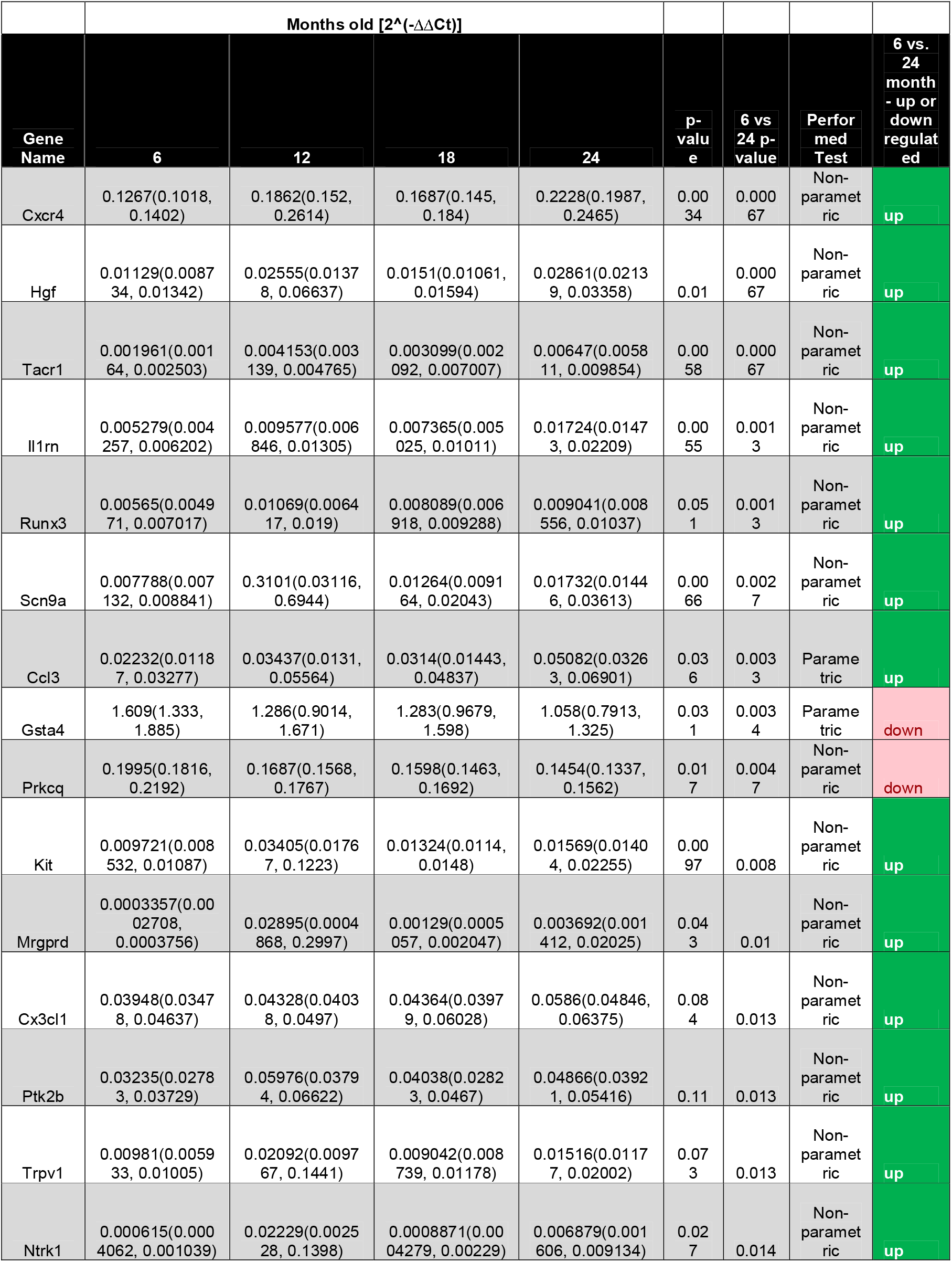
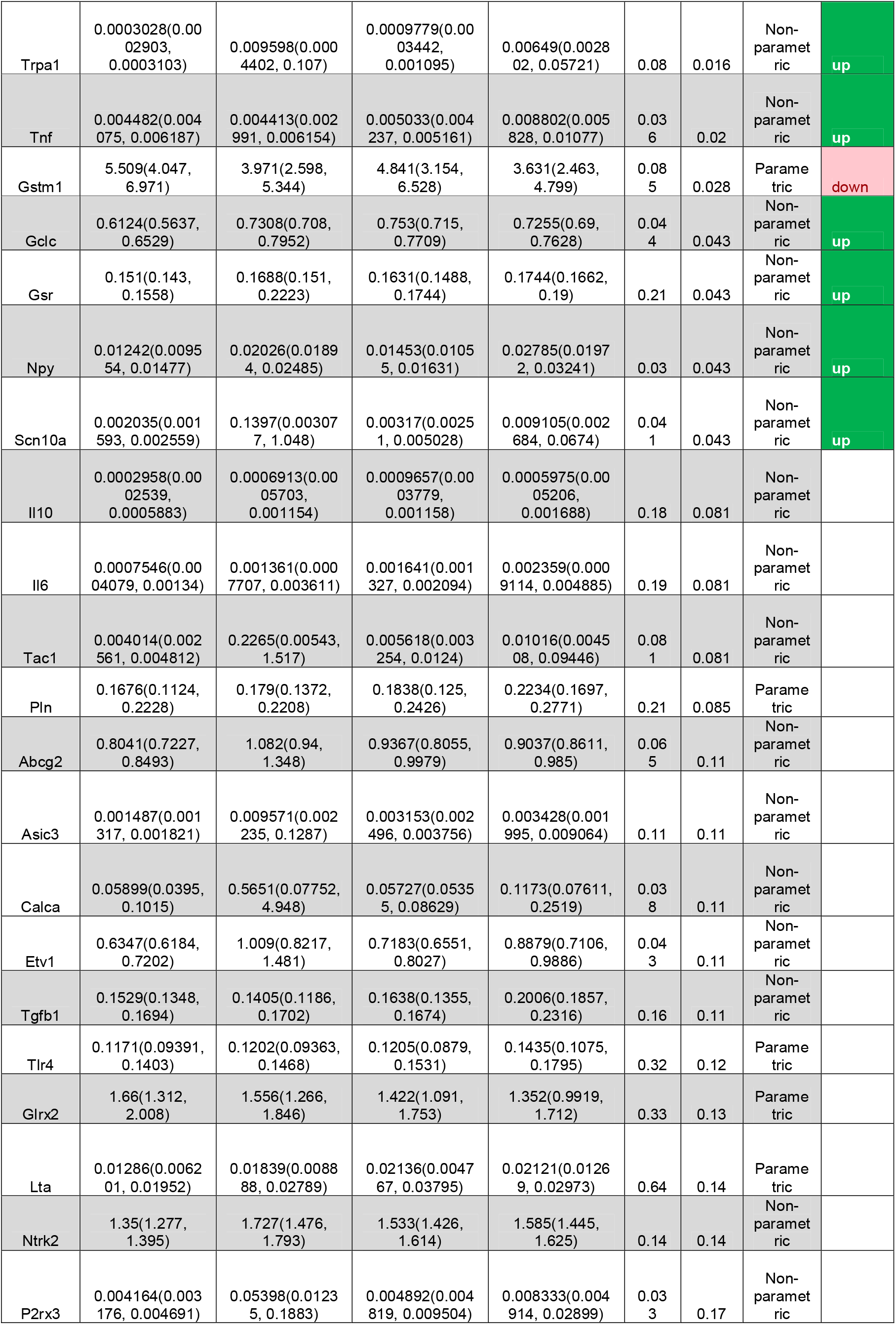
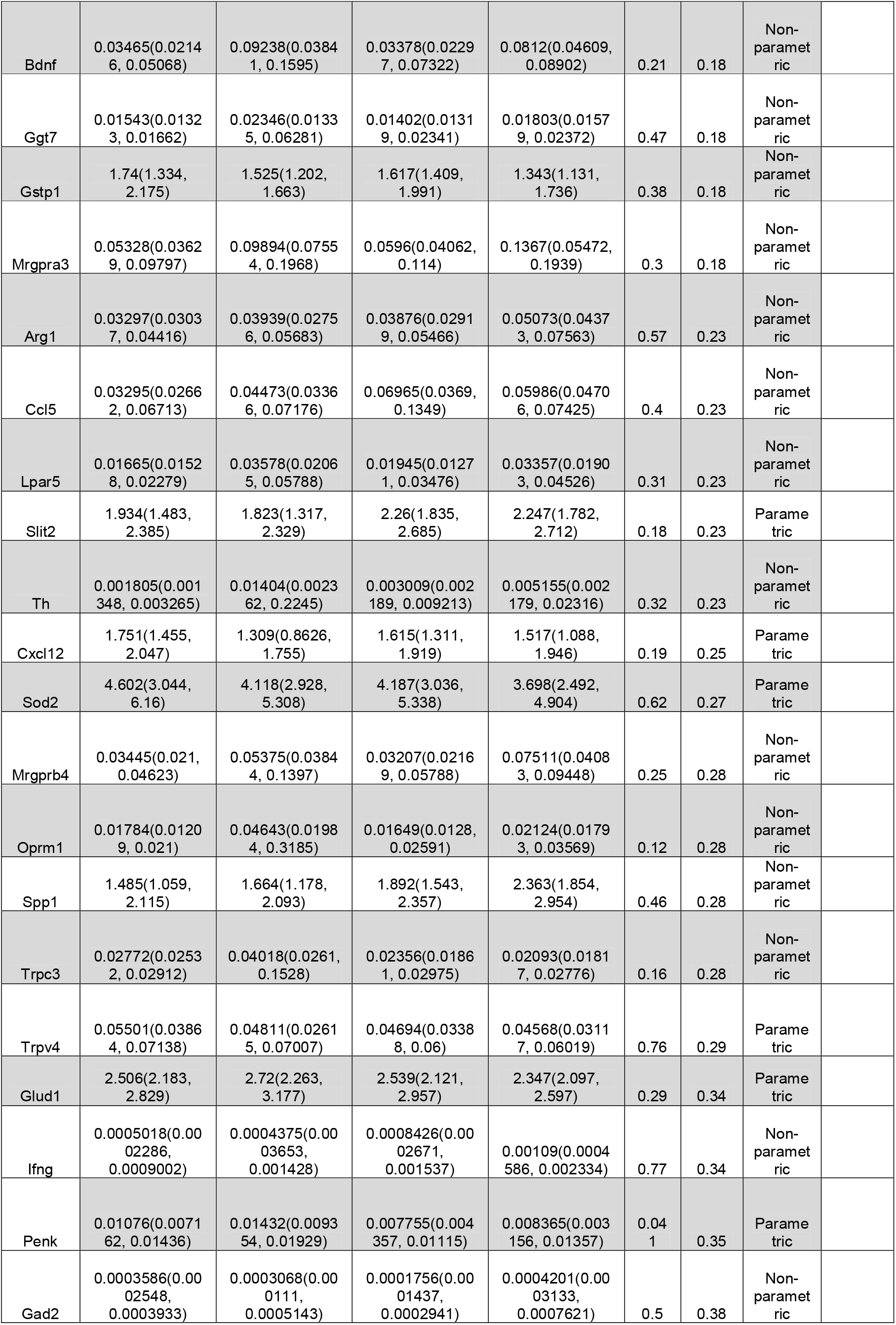
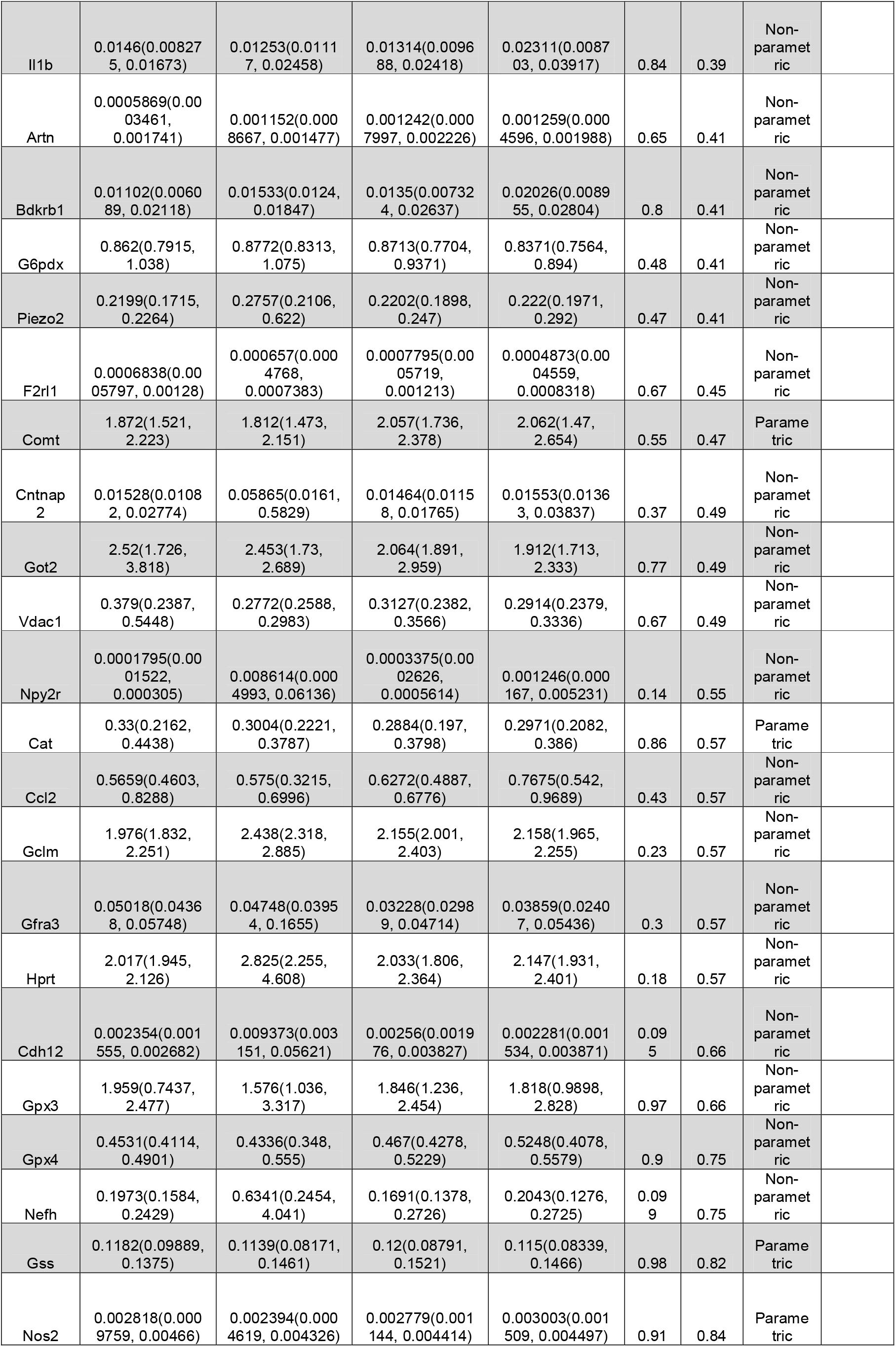
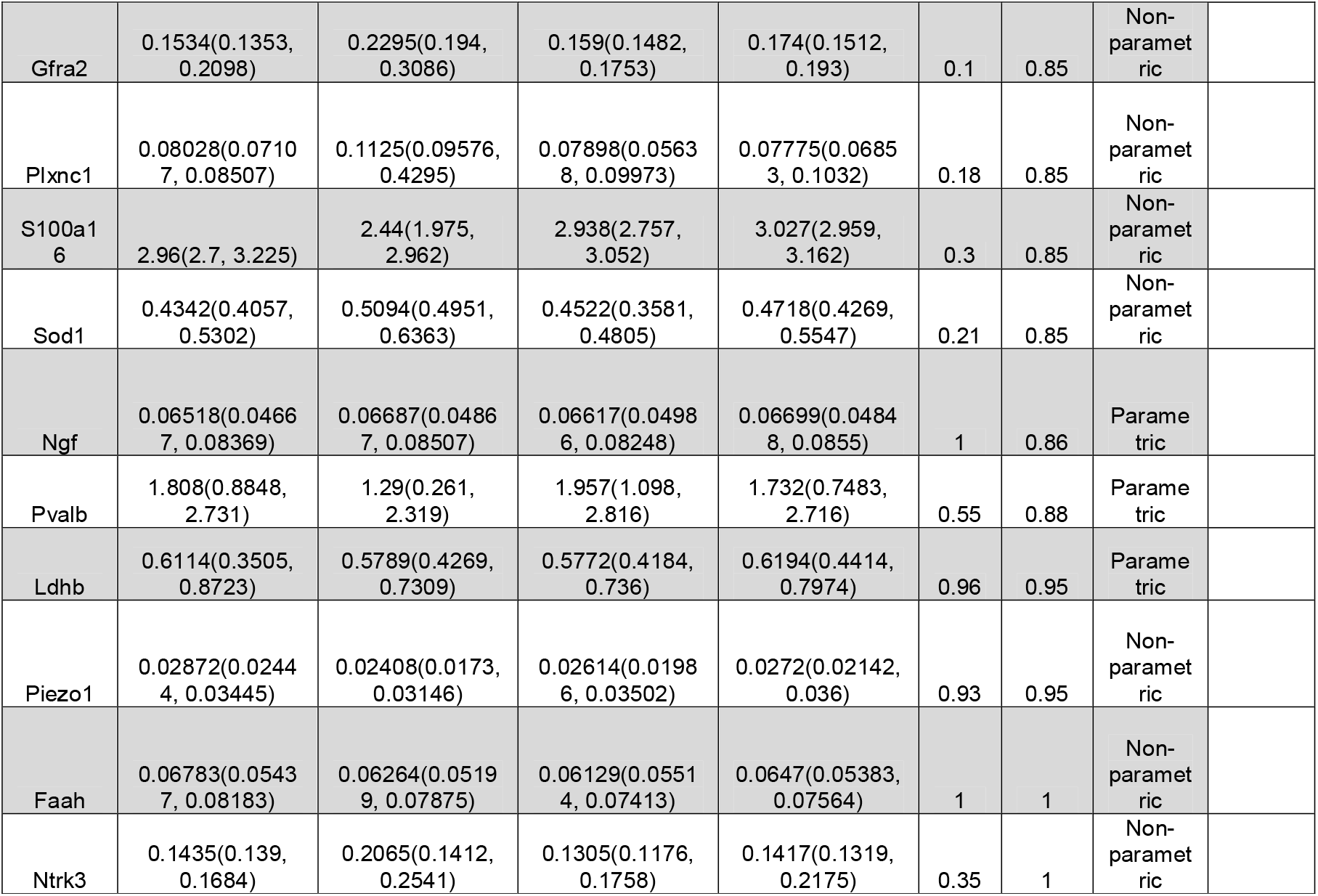
Gene expression in DRGs of female C57BL/6 mice aged 6-months (n=6), 12-months (n=8), 18-months (n=8), and 24-months (n=8) old.

### In situ hybridization of aged human DRGs

To further appreciate the translational relevance of immune-related genes in aged DRGs, we used RNAscope to assess the expression of chemokines *CCL2* and *CCL3* in human DRGs (n=3/sex). We chose these genes because we observed both genes differentially regulated in the mouse DRG gene expression analyses. We assessed 3 male and 3 female human DRGs with an average age at death of 90.9 years and average BMI of 25 (**Table 1**). There was expression of both *CCL2* and *CCL3* in male and female human DRGs (Fig. 6 A, B, D, **and** E). Interestingly, after quantifying this signal in a blinded manner, we observed higher *CCL2+* signal in males compared to females (Fig. 6C), whereas there was higher *CCL3+* signal in females compared to males (Fig. 6F). Together with the mouse data, this suggests a role for chemokine signaling in aging associated DRG changes and perhaps a sexual dimorphism in how these signals change over time.

**Fig. 6.**
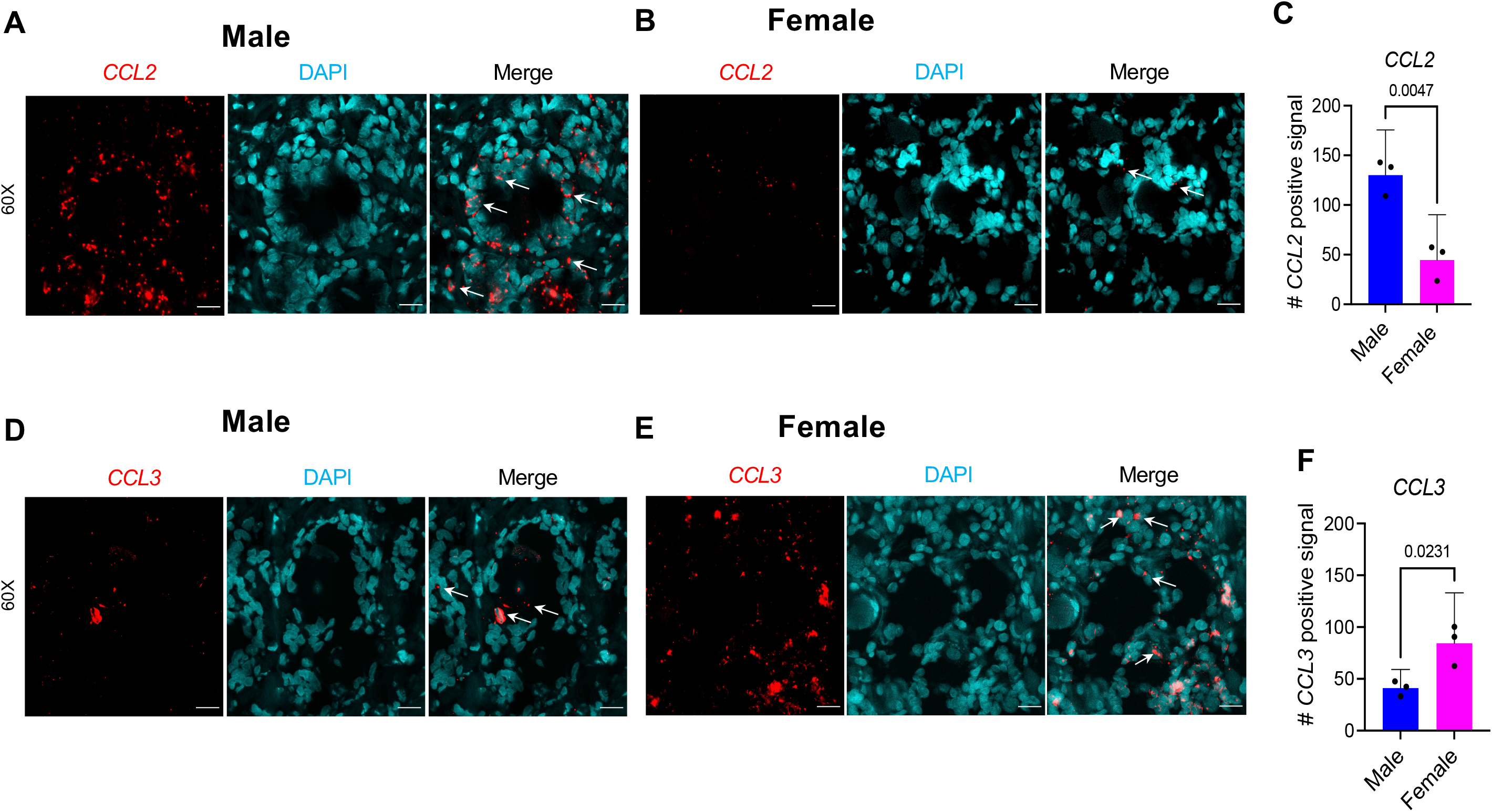
Expression of *CCL2* and *CCL3* in male and female human dorsal root ganglia. (**A-C**) RNAscope used to quantify the expression of *CCL2* or (**D-F**) *CCL3* in serial sections of human dorsal root ganglia (n=6 donors, three male, three female); Mean±SEM. (**A-B, D-E**) Representative sections of (A, D) male and (B, E) female human DRG. 60X magnification (scale bar = 25μm). White arrows indicate examples of positive signal. (**C, F**) RNAscope signal was manually counted in 5-6 60X images and averaged per donor. \**p* ≤ 0.05, \*\**p* ≤ 0.01 by unpaired two-tailed t-test.

## Discussion

Aging is a predominant risk factor for nontraumatic OA development (48). To our knowledge, there have been no studies concurrently evaluating pain behavior, joint damage, and immune cell changes in the DRG in naturally aged mice.

Here, we saw increased signs of OA, including cartilage degeneration, osteophytes, and synovitis in aged male and female mice, with worse OA in males than in females. This supports previous reports that have evaluated naturally occurring mouse OA joint degeneration predominantly in male mice (44, 49–55). McNulty *et al.* reported that older male mice (16-month, 17-month, and 23-month old) had significantly thinner articular cartilage compared to 4.5-month old mice (44). While we did not measure these exact parameters, our results showing significant cartilage degeneration in older mice are consistent with this previous report. Another study showed significant cartilage degeneration in 18- and 25-month old male mice (by safranin-O stain) in the temporomandibular joint (TMJ) (50). In addition, collagen-rich tissues are pre-disposed to common age-related disorders, such as OA, and studies reported age-associated changes in cartilage matrix protein content, such as reduced protein incorporation in all tissues (cartilage, bone, skin), dynamic turnover of matrisome, and reduced collagen synthesis in bone of older mice, compared to younger mice (49, 56). Here, we found mild synovitis and osteophyte presence in both male and female 20-month old mice. One previous report showed no synovial hyperplasia and grade 2 and grade 3 osteophytes in aged mice (18-month old male mice) (57). Other studies showed age-related changes in the meniscus (58), and mild increased OARSI score and synovitis score in WT C57BL/J mice that were 15 months old compared to 3-month old male mice (53). Notably, here we found worse histopathologic OA in male knee joints compared to female, which is opposite of the human OA trend, where females experience increased incidence and severity of OA (2).

Results reported here are also consistent with previous studies showing that aging is correlated with a decline in spontaneous locomotion in mice (29) (59). Tran *et al.* observed a decline in spontaneous locomotion shown by slower gait speeds, and shorter durations of locomotion, rearing, climbing, and immobility in older (22 months) compared to younger (3 month) mice for both sexes (59). They also found a greater age-related decline in males than in females with respect to climbing and rearing. Shoji and colleagues reported age-related behavioral changes in a wide range of behaviors from young adulthood to middle age in C57BL/6J male mice. They found decreased locomotor activity and increased anxiety-like behavior from young adult (2-3 month) to middle aged (8-12 month) mice (29). Here, we did not evaluate spontaneous locomotion or anxiety, but we found that mechanical allodynia, knee hyperalgesia, and grip strength worsen with age to similar degrees in both males and females. This result is consistent with a previous report in a partial menisectomy model of OA, where female mice developed pain-like behavior at the same time as male mice despite reduced chondropathy in females compared to males (60).

Other studies have examined the correlation of age with other OA risk factors such as high fat diet. One study evaluated pain behaviors and joint damage in the context of diet-induced obesity. This study reported obesity from a high-fat or Western diet (HFD) resulted in pain behaviors (increased mechanical sensitivity and altered spontaneous locomotion) in 24-week and 40-week old mice compared to 12-week old mice, and these behaviors preceded joint damage (61). Though the authors did not statistically report difference between age groups (only between different diet groups), if one looks at the chow-fed mice from 12-weeks to 40-weeks of age there is a decrease in spontaneous locomotion and increase in osteophytes in these aged mice, which is consistent with our findings. In another obesity-induced OA study, HFD-induced cartilage catabolism in WT, but not *Tlr4* knockout aged female mice (15-month old), and systemic inflammation was not induced by HFD regardless of mouse genotype (62), suggesting an important role for innate immunity in OA development. In contrast to our results, in MIA-induced OA in mice of different ages, weight bearing asymmetry and mechanical sensitivity were mildly attenuated in 22-month old mice compared to 3-month old mice (30). These results were associated with a reduced microglial response in aged MIA mice (30). In addition, the authors attributed the absence of an early phase pain response in old mice as an age-related reduction in the peripheral inflammatory response (30), which is consistent with results shown here where there was a reduction in total peripheral blood CD45+ leukocytes in both male and female aged mice and peripheral blood T cells in aged male mice (Suppl. Fig. 8). The inconsistency between pain-related outcomes despite similar reductions in immune cell populations in these studies may reflect differential neuro-immune responses to toxin-induced OA pain (MIA) *vs*. spontaneous age-related OA pain.

There are many types of immune cells involved in neuroimmune crosstalk and function (63), including dendritic cells, neutrophils, macrophages, mast cells, and T cells. Studies have indicated that macrophages skew toward an inflammatory phenotype as tissues age, which may be a natural part of aging (64–66). In addition, OA has a prolonged inflammatory phase and lack of repair and remodeling, which may contribute to the development of OA pain (67). How the inflammatory phase gets resolved relies on mediators from macrophages and other immune cells, *e.g*., resolvins, cytokines, and chemokines (68). Moreover, our study here is in line with previous findings in other models demonstrating an important role for DRG macrophages in mediating osteoarthritis pain (69) as well as neuropathic pain (23). Raoof and colleagues showed M1-like macrophages accumulate in the DRG, depletion of macrophages intrathecally resolved OA pain in male and female mice, and altering macrophage signaling to an M2-like phenotype using IL-4/IL-10 fusion protein attenuated OA pain in the MIA model (69). In a nerve injury model, Yu *et al* found that macrophages in the DRG, but not the nerve injury site, are needed for maintenance of mechanical hypersensitivity (23). In other animal models of OA, we and others have found a major role for CCR2 in mediating osteoarthritis pain (34, 70–72), an important chemokine receptor for monocyte/macrophage trafficking and signaling.

Interestingly, sexual dimorphism in the age-related decline of several mouse behaviors has been reported (59), and there are known transcriptional and translational sex differences in mouse sensory neurons (73, 74). Thus, here we studied both males and females, and we found a greater increase in DRG macrophages in aged males than aged females and this coincided with a significant decrease in CCR2+ cells in female DRGs (Figs. 3B, 4B, 3I, 4I). Additionally, we found increased gene expression of chemokines, *CCL2* and *CCL3*, in male and female human DRGs respectively, suggesting a clinical relevance for immune cell modification of sensory neurons. The *CCL2/CCR2* axis has an established role in mediating OA pain (34, 71, 75). Interestingly, blockade of *CCL3* has resulted in diminished pain in a model of neuropathic pain (76), and both CCL2 and CCL3 have been shown to regulate pelvic pain (77). Furthermore, we saw significant increases in MHCII+, CX3CR1+, and CD206+ sub-phenotypes of macrophages in male DRGs, while females had no changes in these populations between young and old DRGs, suggesting a more activated macrophage phenotype in males than females. Deletion of macrophage migration inhibitory factor (MIF) provided protection from OA-like changes in the cartilage, bone, and synovium in male mice aged 22-months old (78). Moreover, T cells are thought to play a more important role in female pain development (79). There have been studies of lymphoid compartments in aged mice (47, 80), and here we show results consistent with the literature of decreased peripheral blood CD4+ T cells only in older males (47). Furthermore, we know there is sexual dimorphism in human immune system aging (81). For example, while age-related epigenomic changes in peripheral blood mononuclear cells occur as soon as late-thirties with similar timing between sexes, the second spike is earlier and stronger in men, and genomic differences between sexes increased after age 65 (81).

While this study of aged mouse knee OA, pain-like behaviors, and DRG immune phenotyping elucidates the complex neuro-immune interactions involved in aging-associated OA pain, it comes with some limitations. We were limited in the number of markers we could evaluate via flow cytometry. We sought to understand these data with other data sets, such as gene expression data of mice DRGs from the NIA rodent colony, which showed increased immune cell recruitment chemokines, *e.g., Ccl2* and *Ccl3*. Furthermore, we used human DRGs to further validate these mouse gene expression analyses providing clinical relevance of these findings. Finally, when studying aging there are always confounding variables that may alter interpretation of findings, such as aging-related obesity and other disorders, *e.g.*, cataracts, dermatitis, etc. Mice studied here did not have any outward signs of disease, and thus, other aging-related disorders do not likely substantially confound the interpretation of our findings. Nonetheless, our study should be considered an observation of features in the knee joint and DRG that accompany aging. In addition, while we do not examine senescence in this study, it certainly plays a role in aging and age-related disorders (82, 83). Immune cell senescence in T cells is well known (84) (33). Here we saw a decrease in circulating T cells in older males, but not females, which is also consistent with previous literature (47), and could be due to T cell senescence.

In summary, this study demonstrates that in mice, aging is accompanied with mild knee osteoarthritis, increased pain-related behaviors, and distinct DRG immune phenotypes, with important differences between sexes. Further research is necessary to evaluate the age-dependent consequences of DRG immune cell changes and their relation to OA pathology and pain. Many of the animal models currently used for studying OA are surgically, injury-, or chemically induced, but only a subset of clinical OA has been linked to post-traumatic injury (85). Collectively, findings we report here may lead to development of better targeted therapies and analgesic options for OA pain, especially for the aging population. Future research will aim to better define how neuroimmune communication contributes to OA pain in both mice and humans. To conclude, studying spontaneous OA in aging mice provides valuable insights that will lead to development of better analgesics for OA pain.

## Acknowledgements

We would like to thank the Rush University Flow Cytometry Core and the OMRF Arthritis and Clinical Immunology Cores. We thank the study participants and staff of the Rush Alzheimer’s Disease Center funded through NIH grants P30AG10161, R01AG17917, R01AG24490. ROSMAP resources can be requested at https://www.radc.rush.edu.

## Author Contributions

All authors were involved in drafting or editing the article and were critically important for providing intellectual content. Study conception and design by TG, AMM, REM. Histopathology staining and analysis by AMO, JL, and CRS. Pain-related behavior testing by TG and SI. Flow cytometry staining and analysis performed by TG. Human DRG tissue processing and RNAscope performed by MJW and quantification by TG. Mouse DRG gene expression acquisition and analyses done by EBP and TMG. Statistical analyses by TG and REM.

## Role of Funding Source

This work was supported by National Institutes of Health (NIH)/National Institute of Arthritis and Musculoskeletal and Skin Diseases (NIAMS) Grants. T32AR073157 and F32AR081129 (TG), R01AR077019 (REM), R01AR064251, R01AR060364, P30AR079206 (AMM), R01AR075737 and I01BX004912 from the VA (CRS). Rheumatology Research Foundation Innovative Research Award (AMM and MJW). R01AG049058 from the NIH National Institute on Aging and I01BX004666 and I01BX004882 from the Department of Veterans Affairs (TMG).

## Conflict of Interest

The following authors declare no conflicts of interest: TG, AMO, SI, MJW, JL, EBP, CRS, and TMG. AMM received consulting fees from Asahi Kasei Pharma Corporation, Eli Lilly, Pfizer, and Collegium Pharma. REM serves as an Associate Editor of *Arthritis & Rheumatology*.

---

**Suppl. Fig. 1.**
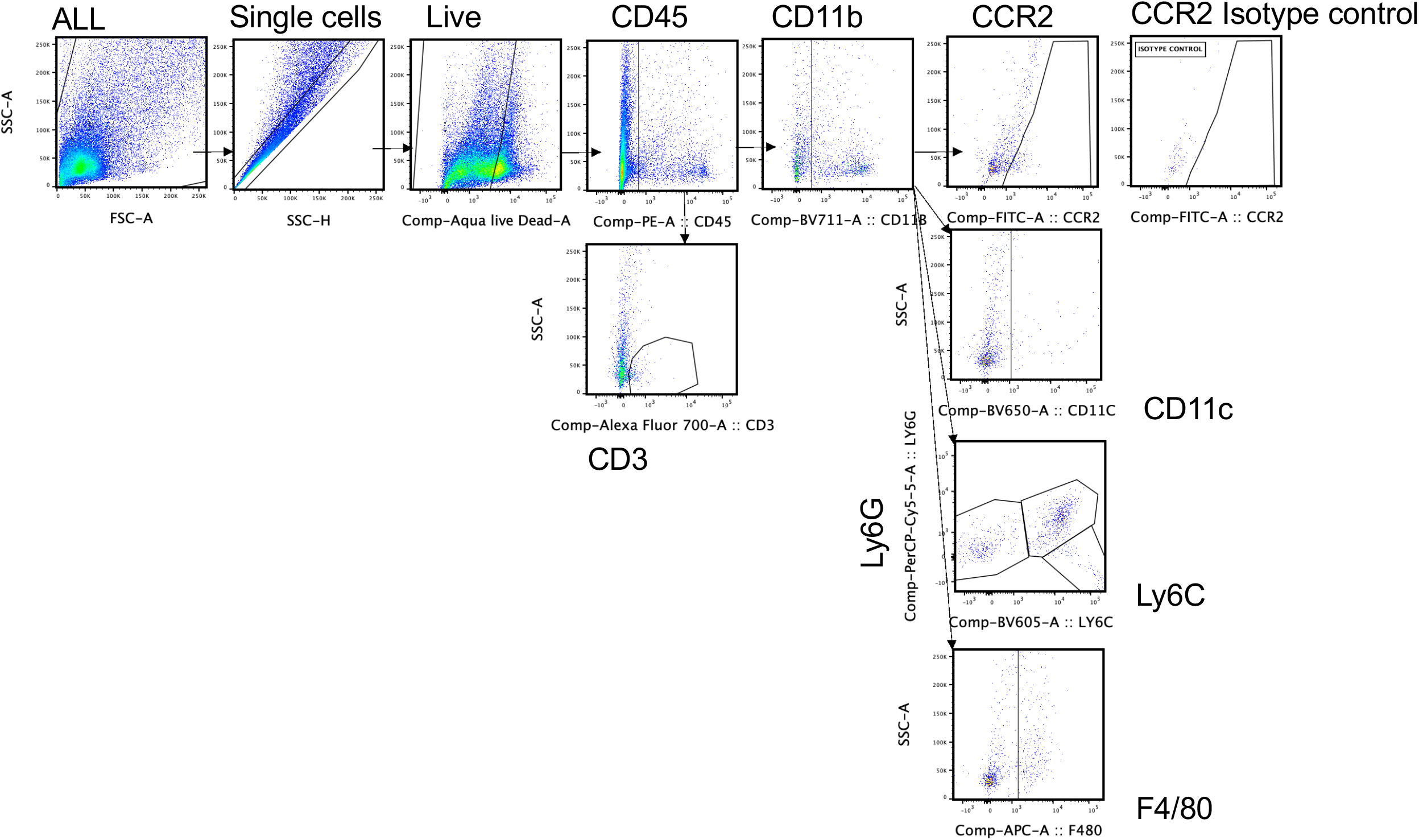
DRG Gating strategy. Representative gating strategy used for all DRG flow cytometry samples in this study. First, cells were gated for single cells to exclude doublets and clumps. Next cells were gated on live cells (Aqua live/dead negative for dye stain). Single live cells were then gated for CD45+ cells, followed by CD11b+ or CD3+ cells. Under CD11b+ cells, remaining poulations were gated for CCR2+ cells, F4/80+ macrophages, CD11c+ dendritic cells, Ly6C+ monocytes, and Ly6G+ granulocytes. Isotype controls were used to draw gates, exemplified here via Isotype control for CCR2.

**Suppl. Fig. 2.**
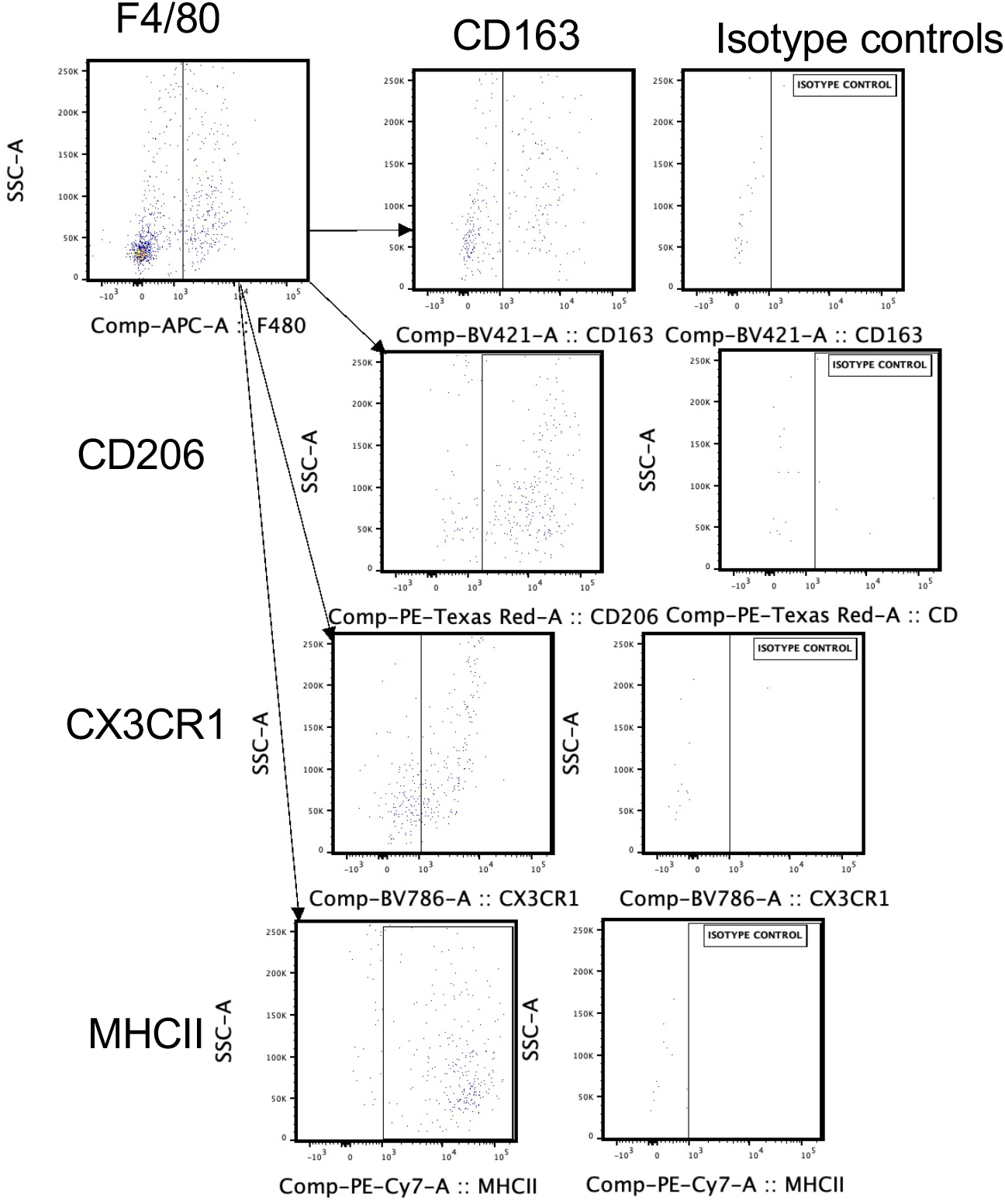
Gating strategy of DRG Macrophage subpopulations. Representative gating strategy used for all DRG flow cytometry samples in this study. Under F4/80+ cell gate, macrophage populations were gated for CD163, CD206, CX3CR1, or, MHCII+ using Isotype controls to inform gating.

**Suppl. Fig. 3.**
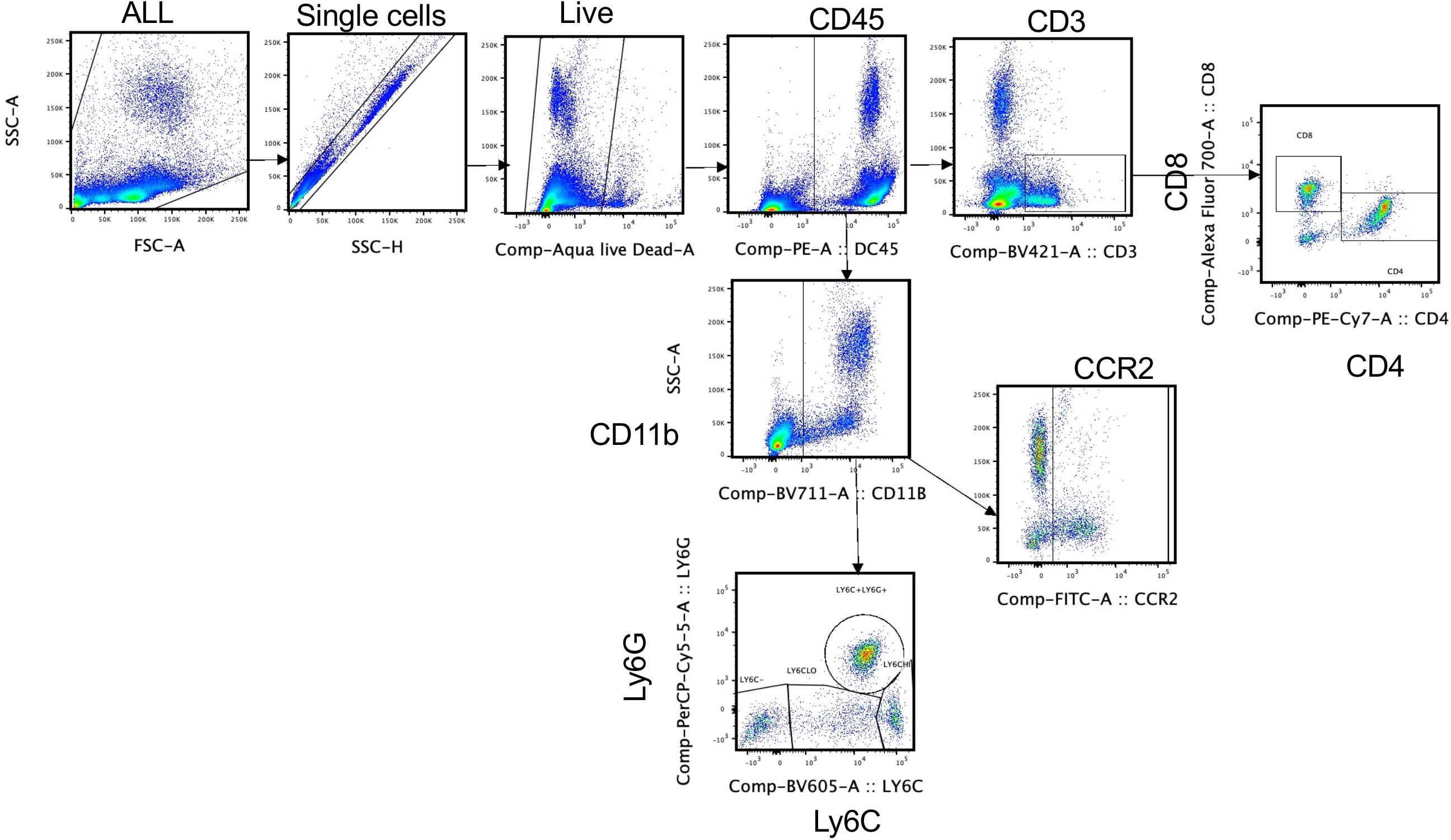
Representative gating strategy used for all peripheral blood flow cytometry samples in this study. First, cells were gated for single cells to exclude doublets and clumps. Next cells were gated on live cells (Aqua live/dead negative for dye stain). Single live cells were then gated for CD45+ cells, followed by CD11b+ or CD3+ cells. Under CD3, cells were gated for CD4 or CD8. Under CD11b+ cells, remaining populations were gated for CCR2+ cells, Ly6C+ monocytes, and Ly6G+ granulocytes.

**Suppl. Fig. 4.**
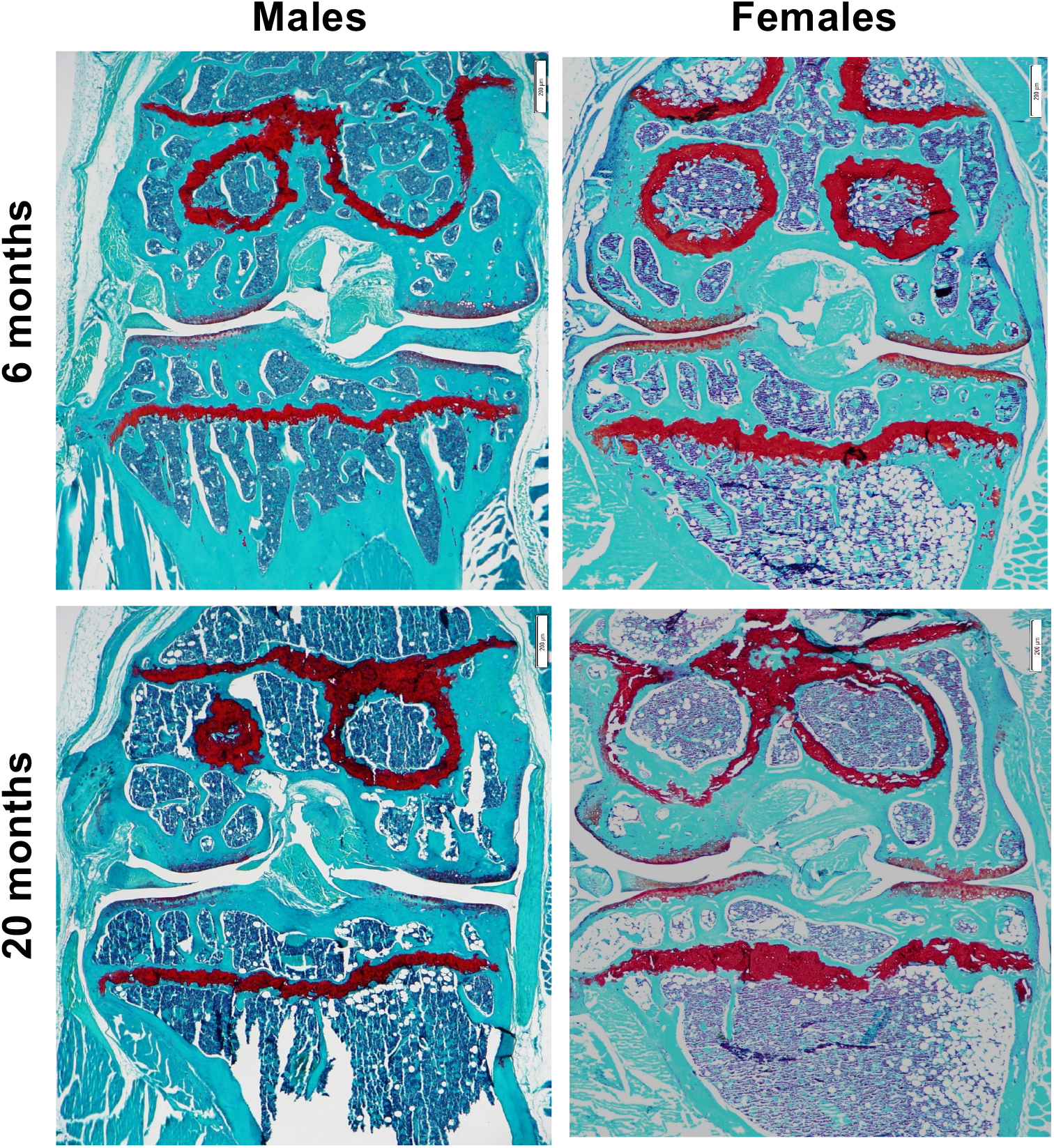
Representative Safranin-O images of aged mouse knee osteoarthritis. Representative images of 6-month or 20-month male or female knee joints stained with Safranin-O. Images taken at 2x magnification.

**Suppl. Fig. 5.**
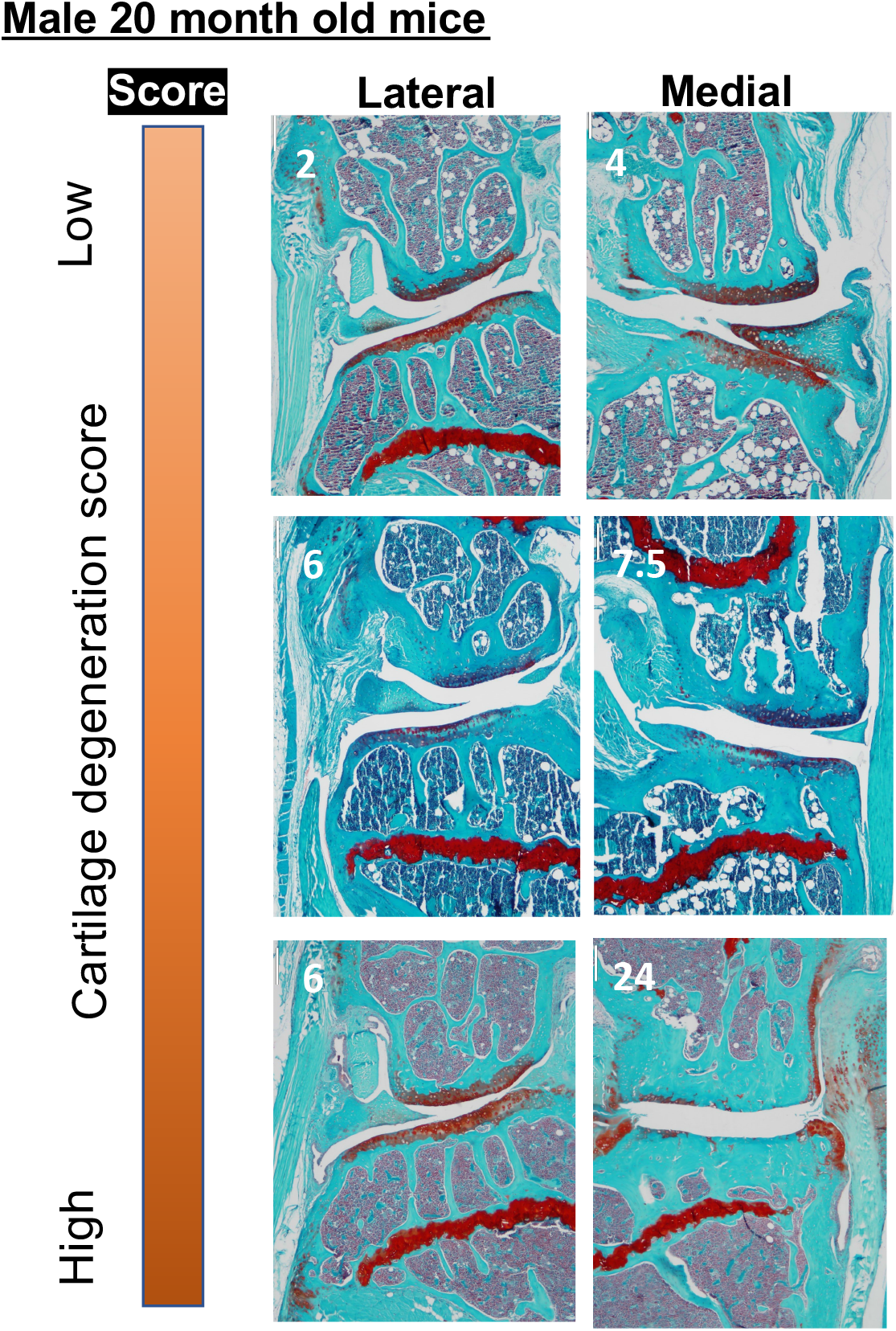
Range of cartilage damage seen in 20 month old male mice. Scores listed in each photo. Representative images of 20-month male knee joints stained with Safranin-O. Images taken at 4x magnification. White numbers on each image show cartilage degeneration score.

**Suppl. Fig. 6.**
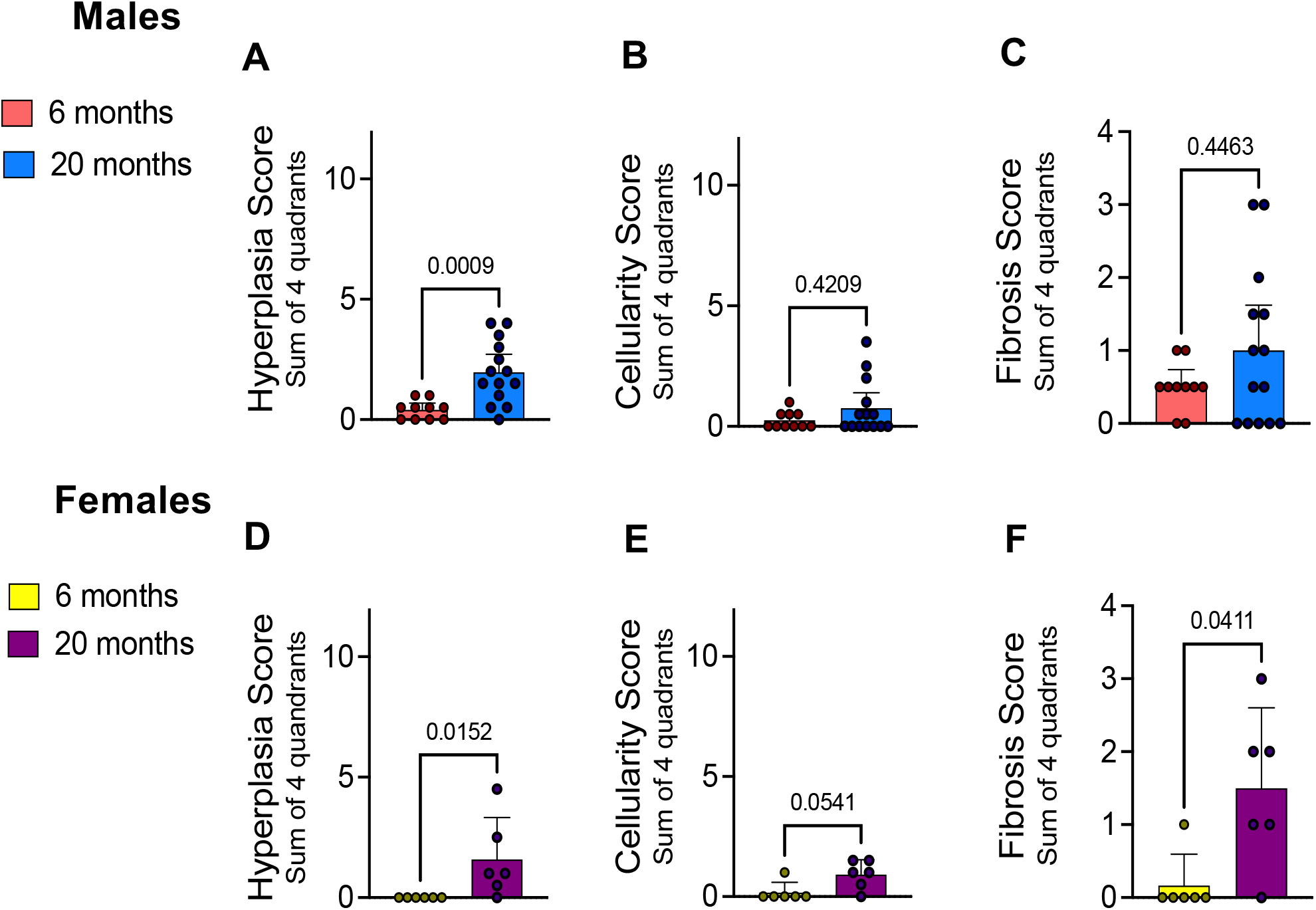
(***A***) Hyperplasia sum score of four tissue quadrants: medial femoral, lateral femoral, medial tibial, and lateral tibial synovium, scored 0-3 per quadrant, maximum sum score of 12 (***B***) Cellularity sum score of four tissue quadrants, scored 0-3 per quadrant, maximum sum score of 12. *(**C**)* Fibrosis sum score of four tissue quadrants, scored 0-1 per quadrant, maximum sum score of 4. *(**A**) – (**C**)* shown for males either 6-month or 20-month old mice. *(**D) – (F**)* Same as in *(A) – (C)* but shown for females aged 6-month or 20-month old mice. Mann-Whitney t-test. Error bars show Mean +/- SEM.

**Suppl. Fig. 7.**
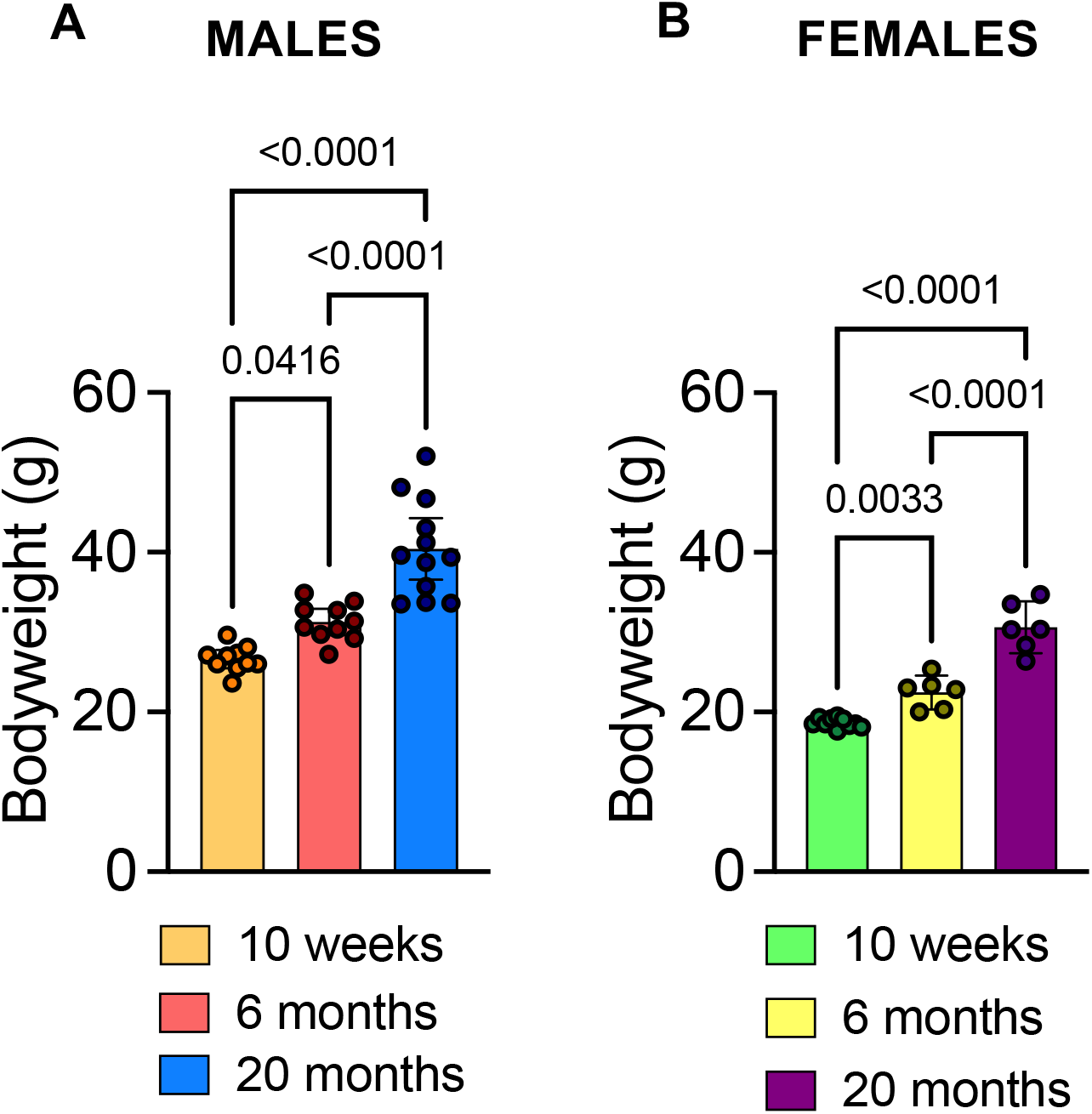
(***A***) Bodyweights of male mice aged 10-weeks, 6-months, or 20-months old in grams. (***B***) yweights of female mice aged 10-weeks, 6-months, or 20-months old in grams. Statistical analysis by ordinary one-way ANOVA. P values stated on graph. Error bars show Mean +/- SEM.

**Suppl. Fig. 8.**
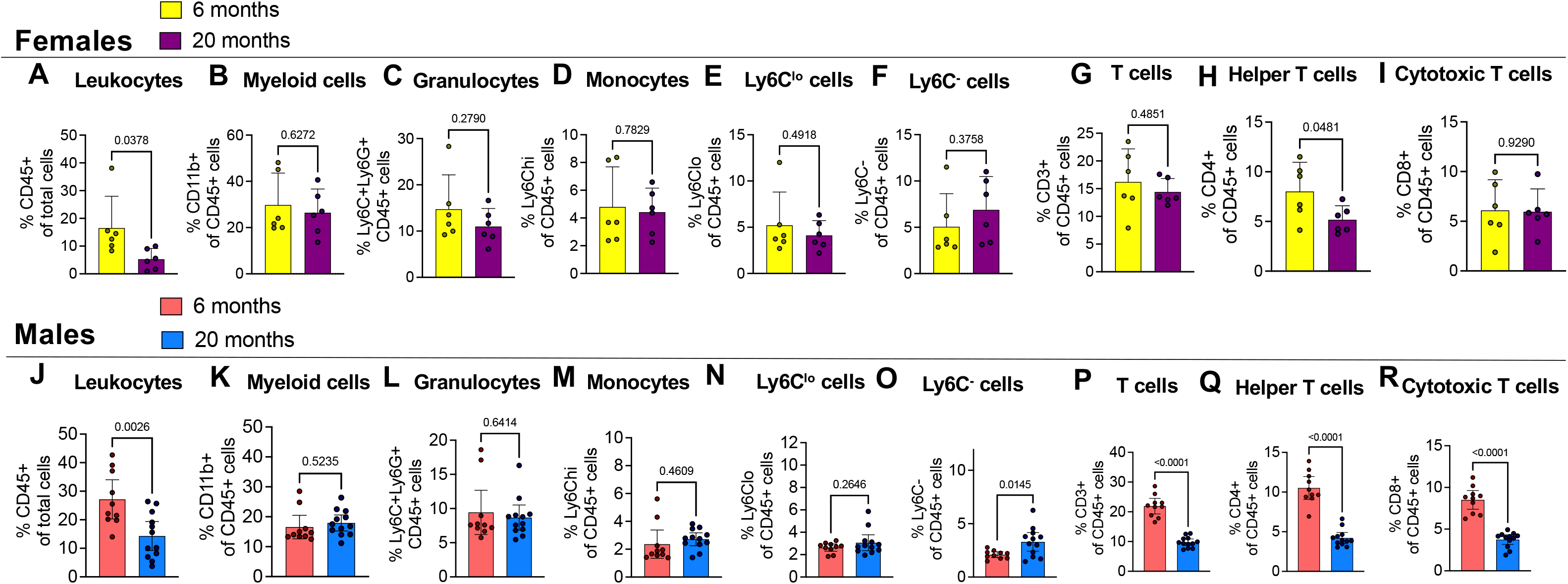
Immune cell populations in peripheral blood of aged male mice. ***(A)*** Frequency of CD45+ leukocytes ***(B)*** CD11b+ myeloid cells ***(C)*** Ly6G+ granulocytes, ***(D)*** Ly6Chi monocytes ***(E)*** Ly6Clo cells ***(F)*** Ly6C-cells, ***(G)*** CD3+ T lymphocytes CD11c+ dendritic cells ***(G)*** Ly6C-antigen presenting precursor cells ***(H)*** CD4+ T helper cells, and ***(I)*** CD8+ cytotoxic T cells in 6-month (n=6), or 20-month (n=6) old female mice.***(J) – (R)*** Same as in ***(A) – (I)*** but for 6- and 2—month old male mice peripheral blood. Flow cytometry gating strategy in Suppl. Fig. 3. Statistical analysis by two-tailed student’s t-test. Significant if p < 0.05. P values stated in graphs. Error bars show Mean +/- SEM.

## Notes

### Competing Interest Statement

The following authors declare no conflicts of interest: TG, AMO, SI, MJW, JL, EBP, CRS, TMG and REM. AMM received consulting fees from Asahi Kasei Pharma Corporation, Eli Lilly, Pfizer, and Collegium Pharma.

